# Disrupting ATXN1 Nuclear Localization in a Knock-in SCA1 Mouse Model Improves a Spectrum of SCA1-Like Phenotypes and their Brain Region Associated Transcriptomic Profiles

**DOI:** 10.1101/2021.12.16.472987

**Authors:** Hillary P. Handler, Lisa Duvick, Jason Mitchell, Marija Cvetanovic, Molly Reighard, Alyssa Soles, Orion Rainwater, Shannah Serres, Tessa Nichols-Meade, Stephanie L. Coffin, Yun You, Brian Ruis, Brennon O’Callaghan, Christine Henzler, Huda Y. Zoghbi, Harry T. Orr

**Affiliations:** Institute of Translational Neuroscience, University of Minnesota, Minneapolis, MN 55455, USA; Department of Neuroscience, University of Minnesota, Minneapolis, MN 55455, USA; Department of Laboratory Medicine and Pathology, University of Minnesota, Minneapolis, MN 55455, USA; Program in Genetics & Genomics and Department of Molecular and Human Genetics, Baylor College of Medicine, Jan and Dan Duncan Neurological Research Institute at Texas Children’s Hospital, Houston, TX 77030, USA; Mouse Genetics Laboratory, University of Minnesota, Minneapolis, MN 55455, USA; Department of Biochemistry, Molecular Biology, and Biophysics, University of Minnesota, Minneapolis, MN 55455, USA; RISS Bioinformatics, Minnesota Supercomputing Institute, University of Minnesota, Minneapolis, MN 55455, USA; Departments of Molecular and Human Genetics, Pediatrics, and Howard Hughes Medical Institute, Baylor College of Medicine, Jan and Dan Duncan Neurological Research Institute at Texas Children’s Hospital, Houston, TX 77030, USA

**Keywords:** Spinocerebellar ataxia type 1, SCA1, neurodegeneration, nuclear localization

## Abstract

Spinocerebellar ataxia type 1 (SCA1) is a dominant trinucleotide repeat neurodegenerative disease characterized by motor dysfunction, cognitive impairment, and premature death. Degeneration of cerebellar Purkinje cells is a frequent and prominent pathological feature of SCA1. We previously showed that transport of ATXN1 to Purkinje cell nuclei is required for pathology, where mutant ATXN1 alters transcription. To examine the role of ATXN1 nuclear localization broadly in SCA1-like disease pathogenesis, CRISPR-Cas9 was used to develop a mouse with the amino acid alteration (K772T) in the nuclear localization sequence of the expanded ATXN1 protein.

Characterization of these mice indicates proper nuclear localization of mutant ATXN1 contributes to many disease-like phenotypes including motor dysfunction, cognitive deficits, and premature lethality. RNA sequencing analysis of genes whose expression was corrected to WT levels in *Atxn1^175QK772T/2Q^* mice indicates that transcriptomic aspects of SCA1 pathogenesis differ between the cerebellum, brainstem, cerebral cortex, hippocampus, and striatum.

## INTRODUCTION

A major challenge facing research on neurodegenerative diseases is the elucidation of key molecular aspects of pathogenesis. While considerable progress has been made in this area for many diseases, the study of pathogenic pathways has typically focused on the molecular aspects of pathogenesis using a limited number of neuronal types – understandably those neurons are characterized as a primary site of pathogenesis. However, such an approach overlooks two critical points; it is becoming increasingly apparent that many neurodegenerative diseases impact multiple cell types and circuits throughout the brain and critical neuronal dysfunction can occur in absence of overt pathology.

Spinocerebellar ataxia type 1 (SCA1) is one of several inherited neurodegenerative diseases caused by expansion of a glutamine-encoding CAG triplet in the affected gene, *ATXN1* in the case of SCA1 (Paulson et al., 2017). SCA1 is characterized by a progressive loss of motor function along with a prominent and consistent degeneration of cerebellar Purkinje neurons. Accordingly, the cerebellum and Purkinje neurons are a major focus of SCA1 pathogenic studies. Our work identified several ATXN1 functional motifs that have a critical role in SCA1 pathogenesis in Purkinje neurons such as the importance of entry of ATXN1 into Purkinje cell nuclei and its interaction with the transcriptional repressor Capicua for Purkinje cell pathogenesis (Klement et al., 1998; Rousseaux et al., 2018). Yet, as SCA1 progresses, symptoms include muscle wasting, cognitive deficits, and bulbar dysfunction resulting in respiratory complications that are a primary cause of premature lethality (Genis et al., 1995; Jacobi et al., 2015; Orengo et al., 2018; Sasaki et al., 1996; Tejwani and Lim, 2020). Correspondingly, a comprehensive pathological study on SCA1 postmortem brains revealed, in addition to cerebellar degeneration, widespread neuronal loss in the primary motor cortex, basal forebrain, thalamus, and brainstem (Rüb et al., 2012). Moreover, longitudinal MRI studies on SCA1 patients identified cerebellar atrophy at diagnosis and changes in brainstem, pons, caudate, and putamen with disease progression (Koscik et al., 2020; Reetz et al., 2013).

As a model resembling the totality of SCA1, *Atxn1^154Q/2Q^* knock-in mice were generated by the insertion of an expanded CAG repeat into one allele of the mouse *Atxn1* gene. These mice express ATXN1[154Q] throughout the brain and display, in addition to ataxia, a range of SCA1-like disease phenotypes including muscle loss, cognitive impairments, and premature lethality (Watase et al., 2002). Here we show that disrupting nuclear localization throughout knock-in mice by replacing the lysine with a threonine at position 772 in the nuclear localization sequence (NLS) of expanded ATXN1, an amino acid change previously shown to block ATXN1[82Q]-induced Purkinje cell disease (Klement et al., 1998), dampens all of the SCA1-like symptoms seen in these knock-in mice. RNA sequencing (RNAseq) was used as means of comparing the disease process in regions of the brain impacted by SCA1. While proper nuclear localization underlies pathogenesis throughout the brain, the RNAseq data indicate that molecular aspects of SCA1 pathogenesis differ among the five regions examined: cerebellum, medulla, cerebral cortex, hippocampus, and striatum.

## RESULTS

### Introduction of a K772T NLS mutation in ATXN1[175Q] does not alter expression of ATXN1

We previously demonstrated that ATXN1 contains a functional NLS in its C-terminus spanning residues 771-774 and that mutating the lysine at position 772 to a threonine disrupted nuclear localization of ATXN1 in cerebellar Purkinje cells. Importantly, when the K772T NLS mutation was introduced into a transgenic mouse model with ATXN1[82Q] expressed exclusively by cerebellar Purkinje cells, the animals no longer developed ataxia or signs of Purkinje cell pathology (Klement et al., 1998). Thus, in this study CRISPR-Cas9 was used to introduce the ATXN1 K772T NLS alteration to the expanded allele of *Atxn1^154Q/2Q^* knock-in mice. To achieve this, three nucleotides in the expanded *Atxn1* gene were altered (Figure 1A). Two changes (AG ➔ CC) were required to generate the K772T amino acid alteration and an additional nucleotide change (C ➔ T) served two purposes: 1) ablation of the PAM site to prevent Cas9 from recognizing and cleaving DNA that had already been altered and 2) introduction of an Alu1 restriction enzyme digest site to expedite genotyping of the mice. After the new mouse line with expanded ATXN1 and the NLS alteration was generated, Sanger sequencing of the *Atxn1* repeat region was performed for both the existing knock-in line and the new NLS mutant knock-in line (Figure 1B). The CAG repeat length was identical in both genotypes, but it expanded from 154 repeats to 175. Accordingly, the knock-in mouse line previously referred to as *Atxn1^154Q/2Q^* is henceforth designated as *Atxn1(175Q)* (*Atxn1^175Q/2Q^*) and the new knock-in model with the ATXN1 K772T NLS alteration is designated as *Atxn1 (175Q)K772T* (*Atxn1^175QK772T/2Q^*).

**Figure 1.**
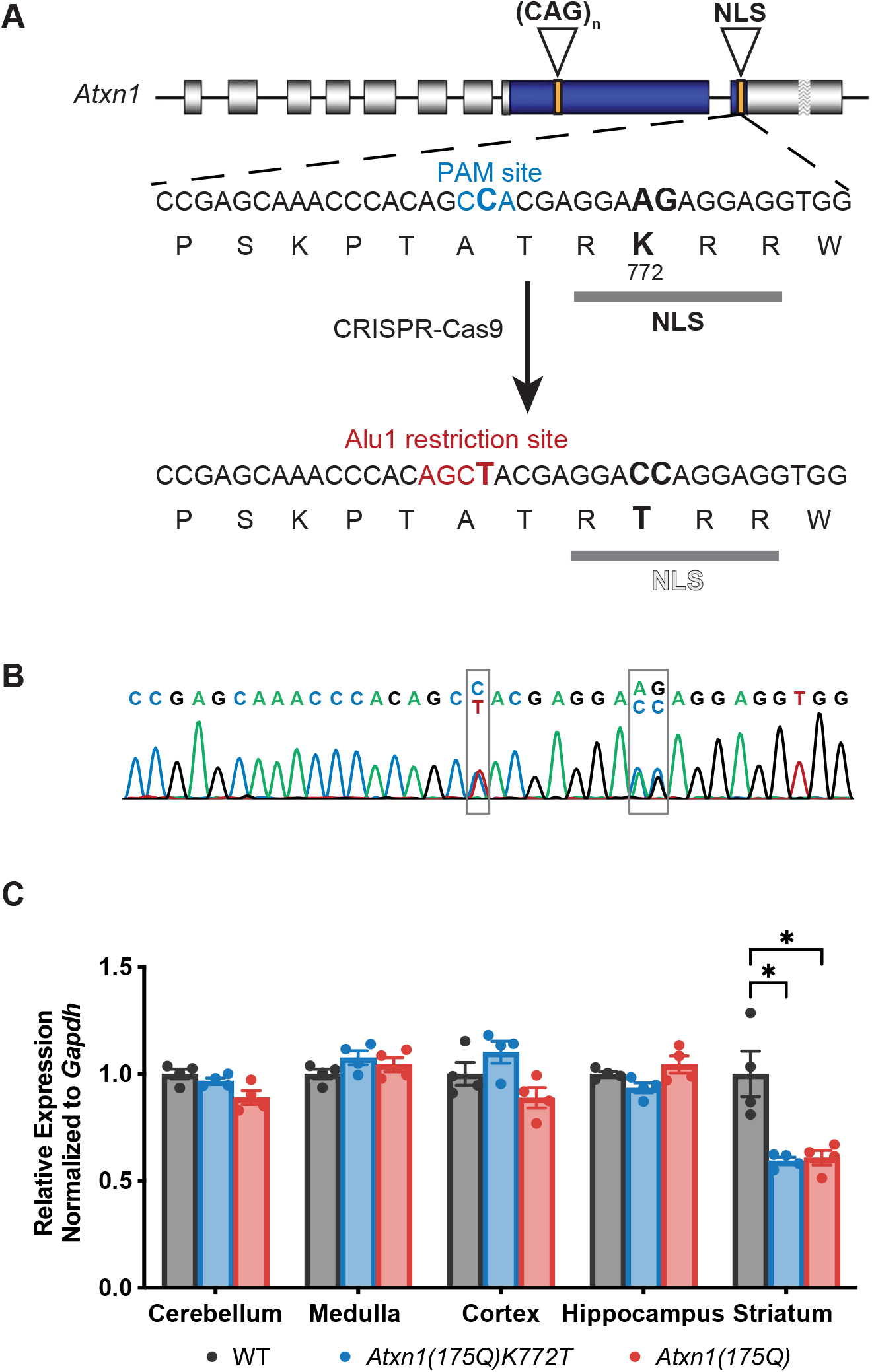
Generation of the *Atxn1^175QK772T/2Q^* mouse model. (A) CRISPR-Cas9 strategy used to create the *Atxn1^175QK772T/2Q^* mouse model. The K772T amino acid alteration in the nuclear localization sequence (NLS) of mouse ATXN1 protein was introduced by changing three nucleotides (bold) in the DNA sequence of the *Atxn1* gene. Two nucleotide alterations (AG ➔ CC) changed a lysine amino acid codon at position 772 of the ATXN1 protein to a threonine amino acid codon (black). Another nucleotide alteration (C ➔ T) ablated the PAM site (blue) and introduced an Alu1 restriction enzyme digest site (red). (B) Sanger sequencing confirming the animals used for breeding were heterozygous for all three desired nucleotide changes. (C) Relative expression of *Atxn1* RNA normalized to *Gapdh* in various brain regions at 26 weeks of age. Data are represented as mean ± SEM. A two-way repeated measure ANOVA with Dunnett’s post hoc test relative to WT expression was performed. Significant results are denoted as * (p<0.05), ** (p<0.01), *** (p<0.001), and **** (p<0.0001). Statistical analysis details can be found in Table S1. Primers used for qPCR can be found in Table S2.

To assess whether the K772T NLS mutation impacted total *Atxn1* RNA expression levels throughout the brain, RT-qPCR was performed in WT (*Atxn1^2Q/2Q^*), *Atxn1(175Q)*, and *Atxn1(175Q)K772T* mice at 26 weeks of age. In the cerebellum, medulla, cortex, and hippocampus, *Atxn1* RNA expression levels were similar between *Atxn1(175Q)K772T* mice and *Atxn1(175Q)* mice and neither were significantly different from WT expression levels. (Figure 1C). Interestingly, while *Atxn1* RNA levels were comparable between *Atxn1(175Q)K772T* mice and *Atxn1(175Q)* mice in the striatum, both genotypes had significantly reduced expression of total *Atxn1* relative to WT mice. Comparing ATXN1[175Q]K772T and ATXN1[175Q] protein expression from *Atxn1(175Q)K772T* mice and *Atxn1(175Q)* mice respectively is complicated by the fact that extractability of expanded ATXN1 from the brain decreases as mice age (Watase et al., 2002). However, using two protein extraction procedures, one to assess subcellular localization and a second to measure protein extractability, we found that the level of ATXN1[175Q]K772T was equal to or higher than the level of ATXN1[175Q] (see below).

### A K772T NLS mutation in ATXN1[175Q] improves a range of SCA1-like disease phenotypes

To establish whether the NLS mutation improves SCA1-like phenotypic deficits historically observed in *Atxn1^154Q/2Q^* mice, we performed a series of assessments comparing WT, *Atxn1(175Q)*, and *Atxn1(175Q)K772T* mice. One prominent SCA1 phenotype is progressive ataxia and motor decline (Zoghbi and Orr, 1995). Motor performance and coordination was assessed by accelerating rotarod in the same cohort of WT, *Atxn1(175Q)*, and *Atxn1(175Q)K772T* mice periodically between 6 weeks of age and 24 weeks of age. On the first trial day with naïve mice at 6 weeks of age, there were no significant differences in performance across the 3 genotypes assessed (Figure 2A). Over the four trial days at 6 weeks of age, the performance of the WT mice improved significantly (p = 0.0008) whereas the performance of the *Atxn1(175Q)* and *Atxn1(175Q)K772T* mice did not change significantly. This indicates that the WT mice were able to learn the motor task and mice with expanded ATXN1 could not. Performance on day 4 from 6 to 24 weeks of age within each genotype did not differ significantly for WT and *Atxn1(175Q)K772T* mice (Figure 2B). In contrast, day 4 performance of the *Atxn1(175Q)* declined significantly between 6 weeks and 24 weeks of age (p < 0.0001). When comparing day 4 performance across genotypes at a given age (Figure 2B), all genotypes were significantly different from all other genotypes at all ages assessed except the comparison between *Atxn1(175Q)* and *Atxn1(175Q)K772T* mice at 6 weeks of age. Together, these findings indicate that the NLS mutation in the *Atxn1(175Q)K772T* mice mitigates the decline in motor performance and coordination with age observed in the *Atxn1(175Q)*.

**Figure 2.**
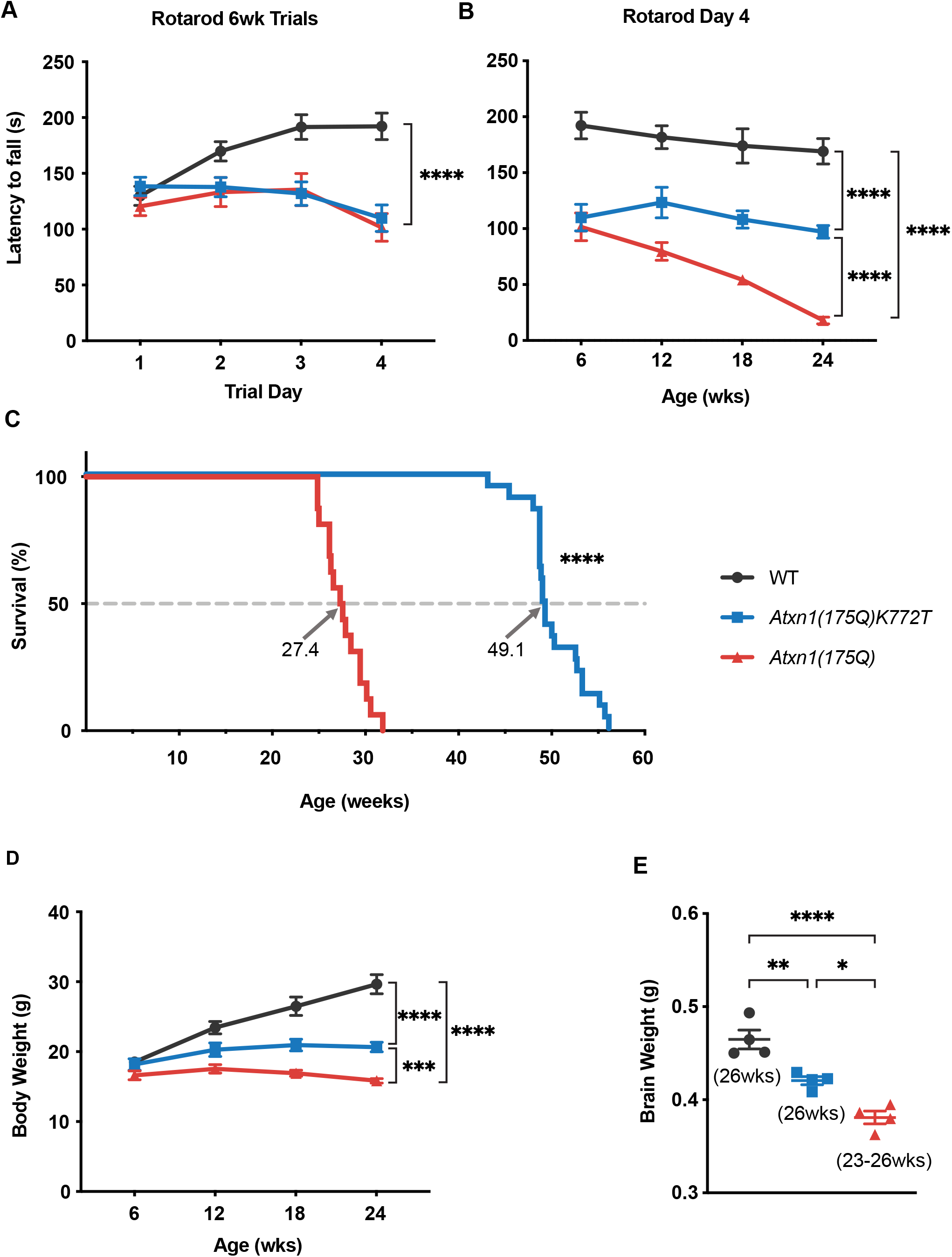
Key SCA1-like phenotypes. (A) Rotarod assessment learning among naïve mice across 4 trial days at 6 weeks of age. (B) Rotarod performance on trial day 4 between 6 and 24 weeks of age. (C) Mouse survival plotted as Kaplan-Meyer curves with median lifespan labeled for each genotype. For *Atxn1(175Q)K772T* n=22 and *Atxn1(175Q)* n= 16. Statistical comparison of survival curves was performed using log-rank Mantel-Cox and Gehan-Breslow-Wilcoxon tests. (D) Body weight measurements between 6 and 24 weeks of age. (E) Brain weight measurements at 23-26 weeks of age. For all genotypes, n=4. Data in (A), (B), and (D) are from the same cohort of mice: WT n=12, *Atxn1(175Q)K772T* n=12, *Atxn1(175Q)* n= 11. Data in (A), (B), (D), and (E) are represented as mean ± SEM. Two-way repeated measures ANOVAs with Tukey’s post hoc test were performed for (A), (B), and (D). One-way ANOVA with Tukey’s post hoc test was performed for (C). Statistical significance is depicted only for genotype comparisons at the last timepoint assessed in (A), (B), and (D). Significant results are denoted as * (p<0.05), ** (p<0.01), *** (p<0.001), and **** (p<0.0001). Additional statistical analyses details can be found in Table S1. *See also Figure S1*

Since cerebellar Purkinje cell degeneration is a well-established cause of motor dysfunction in SCA1, we used RT-qPCR to measure the expression of a group of genes known to be significantly correlated with disease progression in these cells (Ingram et al., 2016). Expression was quantified in WT, *Atxn1(175Q)*, and *Atxn1(175Q)K772T* mice at multiple ages (Figure S1). At 12, 18, and 26 weeks of age, the expression of all 7 genes assessed was significantly decreased in *Atxn1(175Q)* mice relative to WT mice. In comparing *Atxn1(175Q)K772T* to WT mice, only 1 of the 7 genes was significantly different at 12 weeks. By 18 weeks of age, 6 of the 7 genes were significantly different between *Atxn1(175Q)K772T* and WT mice and at 26 weeks, all 7 genes were significantly different. These results show that the NLS mutation delays genetic changes associated with Purkinje cell pathology seen in the *Atxn1(175Q)* mice.

To determine if the NLS mutation improved the premature lethality phenotype observed in *Atxn1^154Q/2Q^* mice (Watase et al., 2002), a survival study was performed on *Atxn1(175Q)* and *Atxn1(175Q)K772T* mice. Median survival of *Atxn1(175Q)* mice was 27.4 weeks, while *Atxn1(175Q)K772T* mice survived significantly longer (p < 0.0001), with a median lifespan of 49.1 weeks (Figure 2C). Thus, the NLS mutation had a dramatic impact on survival. Notably, this relative extension in lifespan is the largest observed resulting from any intervention performed by our group or collaborators (Friedrich et al., 2018; Nitschke et al., 2021; Coffin et al., unpublished data).

*Atxn1^154Q/2Q^* mice also display a failure to gain weight phenotype (Watase et al., 2002). To determine if the NLS mutation improved this phenotype with age, body weight was measured for the same cohort of WT, *Atxn1(175Q)*, and *Atxn1(175Q)K772T* animals periodically between 6 weeks of age and 24 weeks of age. At 6 weeks of age, there were no significant differences in weight between the three genotypes assessed (Figure 2D). At 12 weeks of age, WT mice weighed significantly more than *Atxn1(175Q)* mice (p < 0.0001), but there was not a significant difference in weight between WT and *Atxn1(175Q)K772T* mice nor between *Atxn1(175Q)* and *Atxn1(175Q)K772T* mice. At 18, and 24 weeks of age, however, the average weight for each genotype was significantly different from the other two genotypes. Over time, *Atxn1(175Q)* mice displayed a failure to gain weight phenotype: from 6 weeks to 24 weeks of age, there was not a significant difference in body weight among the *Atxn1(175Q)* animals. Both the WT and *Atxn1(175Q)K772T* mice, however, showed a significant increase (p < 0.0001) in body weight between 6 and 24 weeks of age. Although the weight gain profile of the *Atxn1(175Q)K772T* mice was not as dramatic as that of the WT mice over time, the data indicates that the NLS mutation does substantially improve the failure to gain weight phenotype observed in the *Atxn1(175Q)* mice.

As a measure of neurodegeneration, brain weights were assessed for WT, *Atxn1(175Q)*, and *Atxn1(175Q)K772T* mice. All WT and *Atxn1(175Q)K772T* mice were harvested at 26 weeks of age for brain weight measurements. Two of the *Atxn1(175Q)* mice were sacrificed at 26 weeks of age and the other two *Atxn1(175Q)* mice were designated moribund by veterinary technicians at 23 weeks of age, requiring euthanasia. Brain weight for each genotype was significantly different from the other two genotypes. Accordingly, the NLS mutation significantly mitigates the decreased brain weight observed in *Atxn1(175Q)* mice, but does not restore brain weight of *Atxn1(175Q)K772T* mice to that of WT mice (Figure 2E).

Since cognitive dysfunctions are seen in some SCA1 patients (Bürk et al., 2003) and *Atxn1^154Q/2Q^* mice display cognitive deficits (Asher et al., 2020; Watase et al., 2002), we evaluated the cognitive function of WT, *Atxn1(175Q)*, and *Atxn1(175Q)K772T* mice. The Barnes maze assessment, in which mice are placed on a platform where they use visual cues to locate an escape chamber and exit the arena, was used to compare visuospatial learning, memory, and cognitive strategy in the same cohort of mice at 7 and 17 weeks of age. While mice worked to locate the exit, the Barnes-maze unbiased strategy (BUNS) algorithm (Illouz et al., 2016) was used to classify search strategies and score cognitive performance for each animal (Figure 3A). At 7 weeks of age, WT mice increasingly used the more strategic direct and corrected search methods while both *Atxn1(175Q)K772T* mice and *Atxn1(175Q)* mice were less strategic in their search for the exit (Figure 3B-C). At 17 weeks of age, WT mice had retained the search strategies learned by day 4 of the assessment at 7 weeks of age and maintained use of these strategies across all 4 trial days. Retention of search strategies learned was less robust for *Atxn1(175Q)K772T* mice between the last day of the 7-week assessment and the first trial day at 17 weeks. Notably, at 17 weeks of age, *Atxn1(175Q)K772T* mice increased their use of strategic search methods across the 4 trial days. In contrast, *Atxn1(175Q)* mice failed to retain use of search strategies learned by day 4 of the 7-week assessment and were also unable to improve their use of search strategies across the 4 trial days at 17 weeks (Figure 3D). Clearly, WT mice had the strongest long-term recall of the task and greatest capacity for learning, while *Atxn1(175Q)* mice displayed difficulty learning or remembering the task and their performance declined significantly over time. By the final training day at 17 weeks, both WT and *Atxn1(175Q)K772T* mice had significantly higher cognitive scores than *Atxn1(175Q)* mice (Figure 3E). Together, these results indicate that *Atxn1(175Q)K772T* mice show improved cognition compared to *Atxn1(175Q)* mice and that although *Atxn1(175Q)K772T* mice did not perform as well as WT mice over time, they were more capable of learning and strategy development than *Atxn1(175Q)* mice. Accordingly, the NLS mutation does considerably protect against cognitive decline by the Barnes maze assessment.

**Figure 3.**
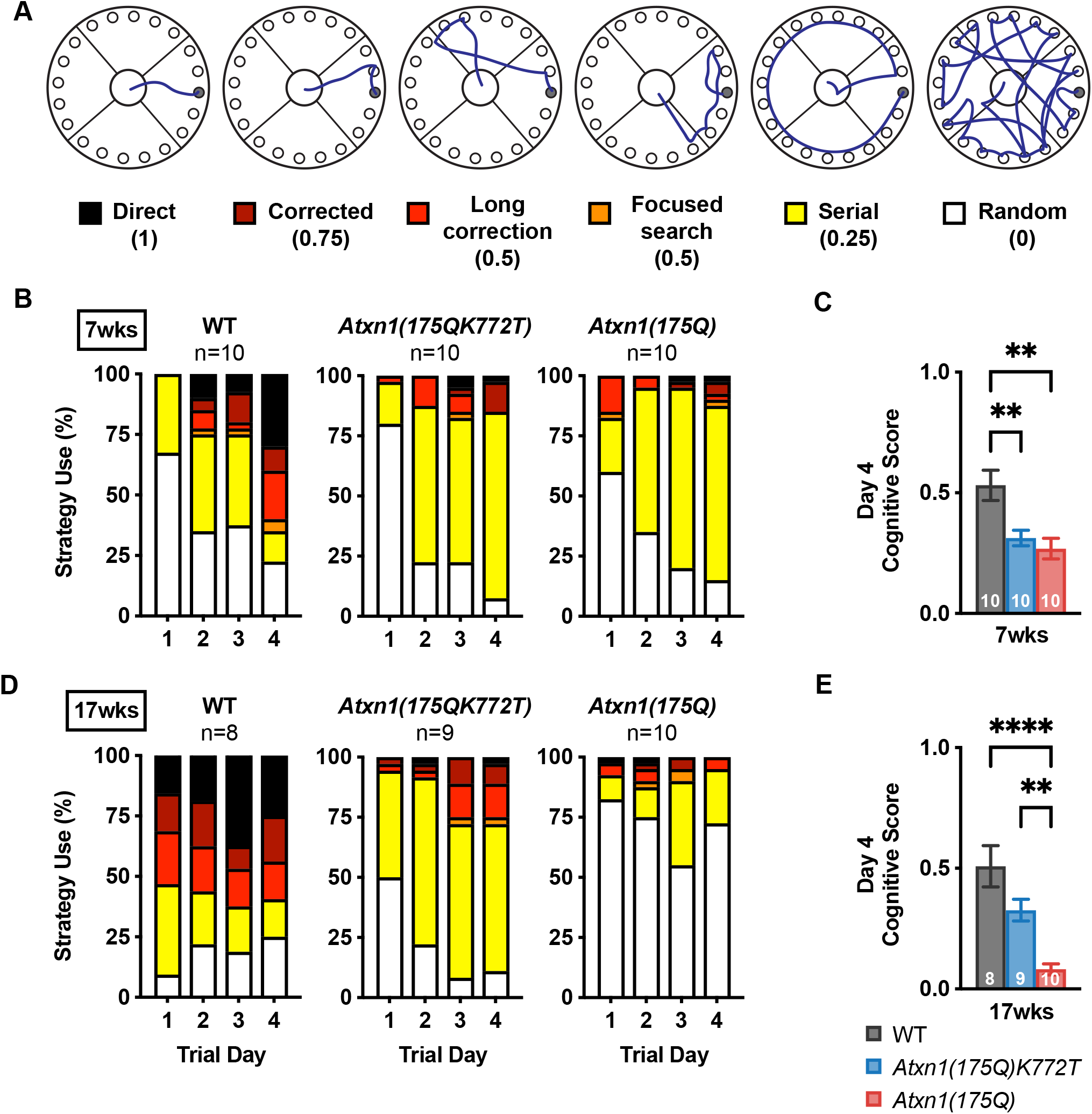
SCA1-like Barnes Maze phenotypes. (A) Representative Barnes Maze search strategy patterns, categorical description, and associated cognitive score. (B) Barnes maze assessment cognitive strategy use percentage among naïve mice across 4 trial days at 7 weeks of age. (C) Barnes maze assessment cognitive scores on trial day 4 at 7 weeks of age. (D) Barnes maze assessment cognitive strategy use percentage across 4 trial days at 17 weeks of age. (E) Barnes maze assessment cognitive scores on trial day 4 at 17 weeks of age. Data in (C) and (E) are represented as mean ± SEM and n values for each genotype are shown in the bottom of each bar. Two-way repeated measures ANOVAs with Tukey’s post hoc test were performed for (B) and (D). One-way ANOVAs with Tukey’s post hoc test were performed for (C) and (E). To assess cognitive score retention between day 4 at 7 weeks and day 1 at 17 weeks for each genotype, paired t-tests were performed for the mice that were tested at both time points. Statistical significance is depicted only for genotype comparisons in (C) and (E). Significant results are denoted as * (p<0.05), ** (p<0.01), *** (p<0.001), and **** (p<0.0001). Additional statistical analyses can be found in Table S1. *See also Figure S2*

Another behavioral measure of cognitive performance used was the contextual fear conditioning assay. Mice were given a series of 5 foot shocks in an environment with unique visual and olfactory cues. After a 24-hour period in their home cage, the mice were placed into the same environment where shocks were administered and the amount of time spent freezing was measured as an indication of fear associated with the expectation of receiving a foot shock. This contextual fear conditioning assessment was performed with WT, *Atxn1(175Q)*, and *Atxn1(175Q)K772T* mice at 8 weeks of age (Figure S2). There was no significant difference in fear response between the WT and *Atxn1(175Q)K772T* mice. Both the WT and *Atxn1(175Q)K772T* mice spent a significantly larger proportion of their time freezing than the *Atxn1(175Q)* mice. This result further supports the idea that the NLS mutation is protective against the cognitive deficits in associative memory observed in *Atxn1(175Q)* mice at 8 weeks of age.

### The K772T ATXN1[175Q] NLS mutation reduces nuclear localization, improves solubility, and alters formation of nuclear inclusions of expanded ATXN1 throughout the brain

We evaluated the impact of the NLS mutation on nuclear entry of ATXN1[175Q] throughout the brain by assessing the subcellular localization, extractability, and nuclear inclusion formation of ATXN1[175Q] and ATXN1[175Q]K772T from *Atxn1(175Q)* mice and *Atxn1(175Q)K772T* mice respectively. To quantify the relative proportions of ATXN1 in the nucleus and cytoplasm, subcellular fractionation was performed on cerebellum, medulla, cerebral cortex, hippocampus, and striatum tissue (Figure 4 and Figure S3). Of note, the proportion of ATXN1[2Q] in the nucleus was constant across brain regions, age, and genotype, averaging 0.87 (range 0.75-0.93). In comparing *Atxn1(175Q)* and *Atxn1(175Q)K772T* mice at 5 weeks of age, there was a significantly higher proportion of ATXN1[175Q] in the nucleus than ATXN1[175Q]K772T for all brain regions assessed. Since quantification of expanded ATXN1 protein quickly becomes less reliable with age (Watase et al., 2002; Figure 5 and Figure S4), subcellular fractionation was not performed on *Atxn1(175Q)* animals at a later age. In the *Atxn1(175Q)K772T* mice, where the extractability of expanded ATXN1 is much improved (Figure 5 and Figure S4), there was no significant change in the nuclear proportion of ATXN1[175Q]K772T between 5 weeks and 42 weeks of age for the medulla, cortex, striatum, and hippocampus (Figure 4). Overall, these findings show that the NLS mutation, while not completely preventing nuclear entry, significantly reduces nuclear localization of expanded ATXN1 throughout the brain and across the lifespan of *Atxn1(175Q)K772T* mice.

**Figure 4.**
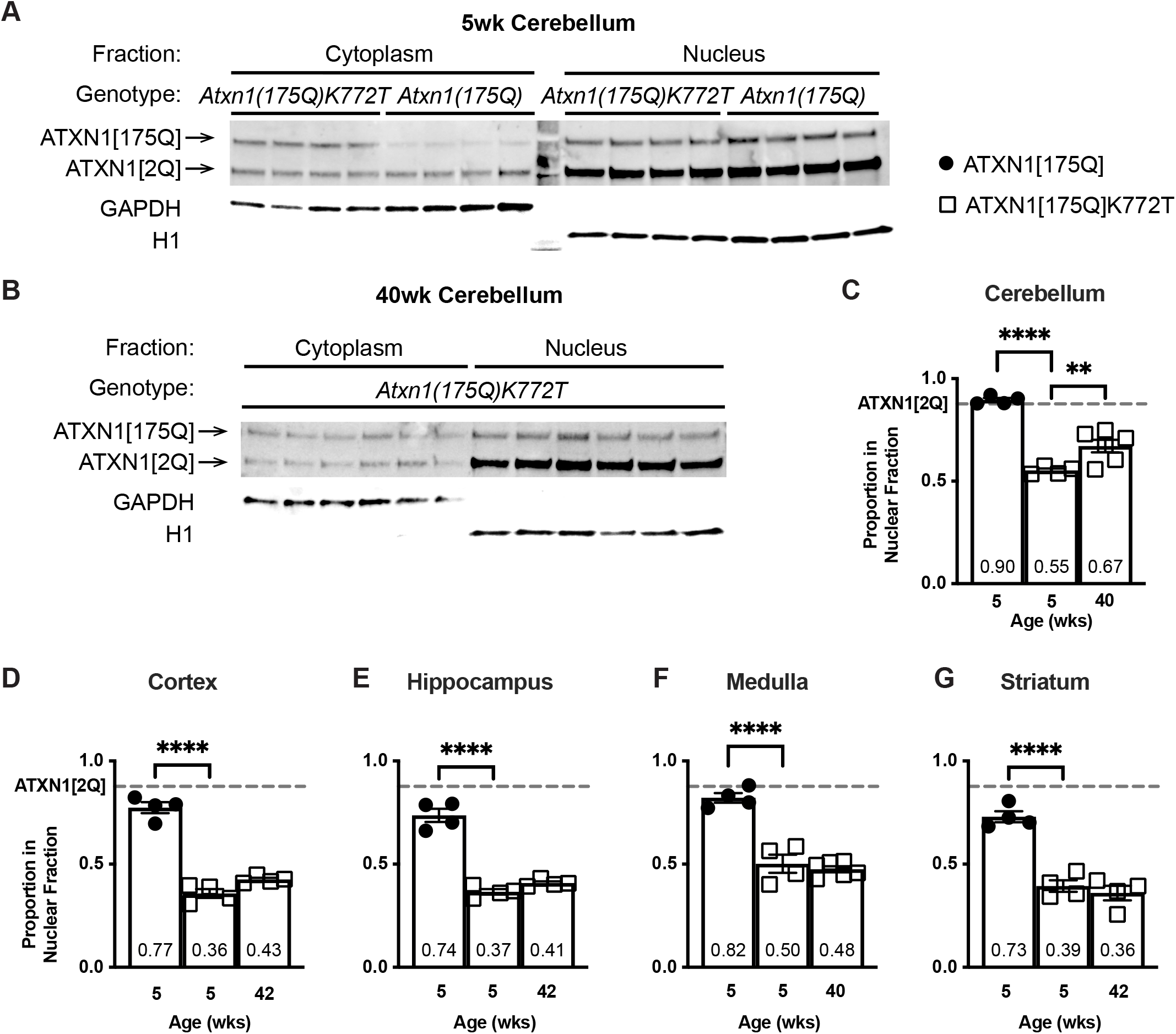
Subcellular fractionation. (A and B) Subcellular fractionation Western blot used to quantify nuclear proportion of expanded ATXN1 from cerebellar lysates in *Atxn1(175Q)* and *Atxn1(175Q)K772T* mice at 5 weeks of age (A) and in *Atxn1(175Q)K772T* mice at 40 weeks of age (B). GAPDH was used as a cytoplasmic marker and H1 was used as a nuclear marker to confirm purity of subcellular fractions. n=4 mice per genotype at 5 weeks of age (A) and n=6 mice at 40 weeks of age (B). (C-G) Proportion of expanded ATXN1 in the nucleus of cells from the cerebellum (C), cerebral cortex (D), hippocampus (E), medulla (F), and striatum (G) at 5 weeks for *Atxn1(175Q)* and *Atxn1(175Q)K772T* mice and at 40-42 weeks for *Atxn1(175Q)K772T* mice. For all brain regions, n=4 mice per genotype at 5 weeks of age and n=4-6 mice at 40-42 weeks of age. Nuclear proportion of ATXN1 was determined by dividing the intensity of nuclear expanded ATXN1 bands by the intensity of expanded ATXN1 bands in the nucleus and cytoplasm combined for a given genotype at a given time. The dashed line represents the average nuclear proportion of ATXN1[2Q] protein product across the three assessments for a particular brain region. Data are represented as mean ± SEM. One-way ANOVAs with Dunnett’s post hoc test relative to the nuclear proportion of ATXN1[175QK772T] at 5 weeks were performed. Significant results are denoted as * (p<0.05), ** (p<0.01), *** (p<0.001), and **** (p<0.0001). Statistical analysis details can be found in Table S1. *See also Figure S3 and S5*

**Figure 5.**
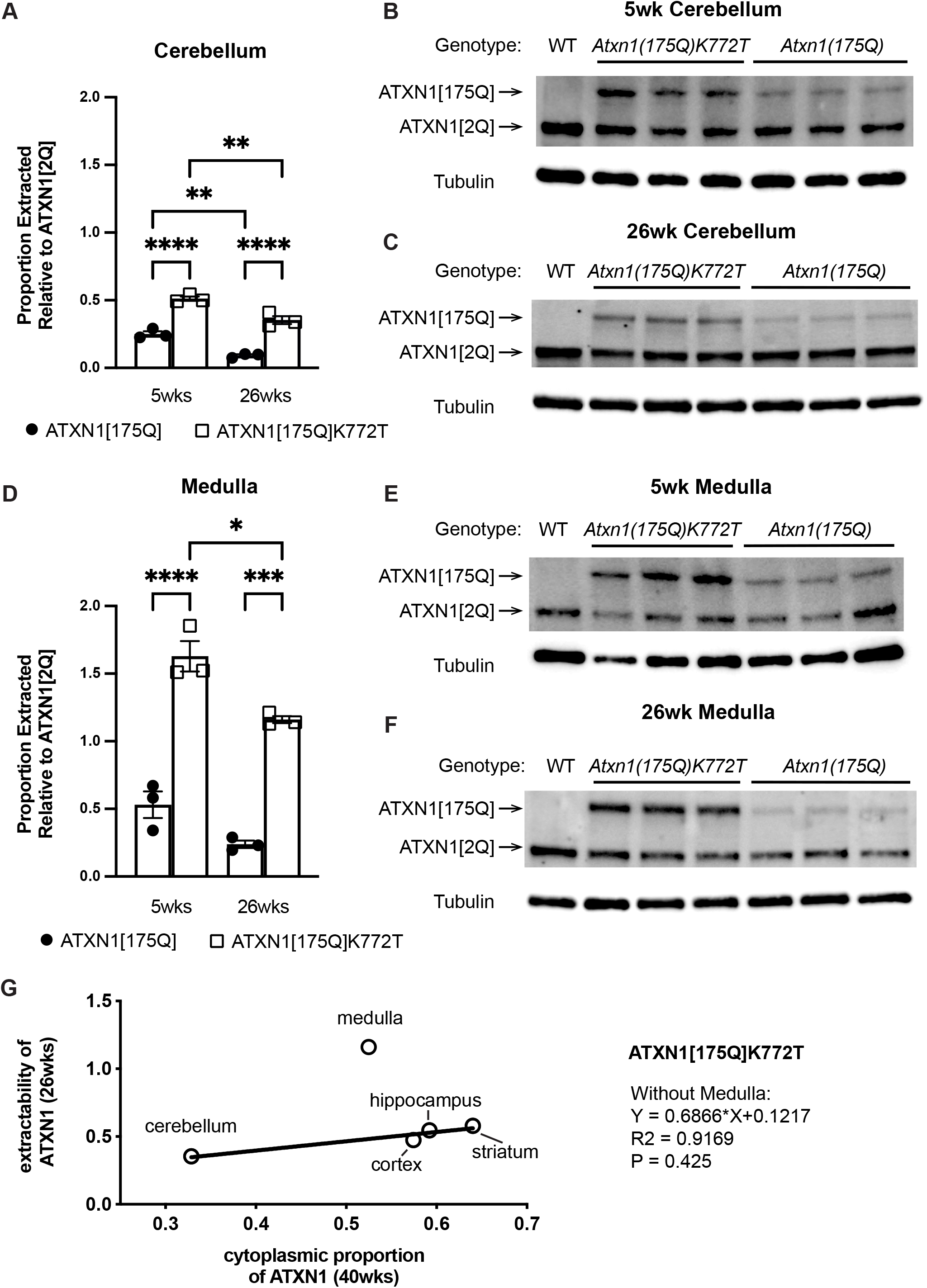
ATXN1 extractability. (A and D) Cerebellum (A) and medulla (D) extractability of expanded ATXN1 protein products relative to ATXN1[2Q] in *Atxn1(175Q)* and *Atxn1(175Q)K772T* mice at 5 and 26 weeks of age. Extractability was determined for each animal by dividing intensity of the expanded ATXN1 band by the intensity of the ATXN1[2Q] band. Data are represented as mean ± SEM and n=3 mice per genotype at each age. Two-way repeated measures ANOVAs with Tukey’s post hoc test were performed. Significant results are denoted as * (p<0.05), ** (p<0.01), *** (p<0.001), and **** (p<0.0001). Statistical analysis details can be found in Table S1. (B and C) Western blots used for extractability quantification of expanded ATXN1 protein products from cerebellar lysates in *Atxn1(175Q)* and *Atxn1(175Q)K772T* mice at 5 weeks of age (B) and 26 weeks of age (C). (E and F) Western blots used for extractability quantification of expanded ATXN1 protein products from medulla lysates in *Atxn1(175Q)* and *Atxn1(175Q)K772T* mice at 5 weeks of age (E) and 26 weeks of age (F). (G) Relationship between extractability of ATXN1[175Q]K772T and proportion of ATXN1[175Q]K772T]found in the cytoplasm for all brain regions assessed. Simple linear regression was performed excluding the medulla data. *See also Figure S4 and S5*

As noted above, the extractability of ATXN1[154Q] decreased rapidly with age, i.e. between 2 and 9 weeks of age in *Atxn1^154Q/2Q^* mice (Watase et al., 2002). In whole-region extracts from the cerebellum, medulla (Figure 5), cortex, hippocampus, and striatum (Figure S4), ATXN1[175Q]K772T was significantly more extractable than ATXN1[175Q] (relative to ATXN1[2Q]) at both 5 weeks and 26 weeks of age. From 5 weeks to 26 weeks, extractability of ATXN1[175Q] did not change significantly in any brain region except the cerebellum, in which extractability decreased over time. While the extractability of ATXN1[175Q]K772T was much greater than the extractability of ATXN1[175Q] at both time points, ATXN1[175Q]K772T did become less extractable over time in all brain regions assessed. For both the extractability blots and the subcellular fractionation blots at 5 weeks of age, the total amount of ATXN1[175Q]K772T was greater than or equal to the level of ATXN1[175Q] in all brain regions assessed (Figure S5). This indicates that, although quantification of expanded ATXN1 is complicated by a reduction in extractability with age, the *Atxn1(175Q)K772T* mice express mutant protein at levels at or above the levels expressed in *Atxn1(175Q)* mice.

Figure 5G depicts the relationship between the extractability of ATXN1[175Q]K772T and the proportion of ATXN1[175Q]K772T found in the cytoplasm for the different brain regions assessed. With the exception of the medulla, there was a strong positive correlation between extractability and cytoplasmic proportion of ATXN1[175Q]K772T. The extractability results seen in the medulla of the *Atxn1(175Q)K772T* mice were a clear outlier compared to the other brain regions as ATXN1[175Q]K772T was more extractable than the WT protein product, ATXN1[2Q], in this region.

Once in the nucleus, expanded ATXN1 accumulates into punctate inclusions (Irwin et al., 2005). As an additional means of assessing the effect of the NLS mutation on nuclear localization of ATXN1[175Q], we quantified ATXN1 nuclear inclusions in brain regions associated with SCA1-like phenotypes. Perfused brain tissue from *Atxn1(175Q)* and *Atxn1(175Q)K772T* mice was immunofluorescently labeled for ATXN1 and NUP62, a nuclear membrane marker (Figure 6A, 6D). An Imaris pipeline was developed to generate a nuclear mask using the NUP62 staining, remove all fluorescent ATXN1 signal outside of the nuclear mask (Figure 6B, 6E), and analyze only ATXN1 staining within the cellular nucleus (under the mask) (Figure 6C, 6F). In each brain region, the percentage of all cells analyzed with at least one nuclear inclusion was recorded for each mouse. Since cerebellar Purkinje cells do not develop nuclear inclusions until relatively late in disease progression (Watase et al., 2002), this cell population was analyzed in *Atxn1(175Q)* and *Atxn1(175Q)K772T* mice at 21 weeks of age. No Purkinje cell inclusions were observed at this age for the *Atxn1(175Q)K772T* mice while over half of the *Atxn1(175Q)* Purkinje cells had at least 1 inclusion, which amounted to a significant difference (p < 0.0001) in inclusion frequency between the two genotypes (Figure 6G). Nuclear inclusions were assessed in ventral medulla (Figure 6H and Figure S6), cerebral cortex (Figure 6L), CA1 of the hippocampus (Figure 6M), and dorsal striatum (Figure 6N) at 12 weeks of age because inclusions are typically present in *Atxn1^154Q/2Q^* mice by this time point in these regions. At this age, the medulla was the only brain region with a significant difference (p = 0.0456) in number of cells with a nuclear inclusion between *Atxn1(175Q)* and *Atxn1(175Q)K772T* mice. There were no inclusions detected in any of the medulla cells in *Atxn1(175Q)K772T* mice and there was a large range in the number of medulla cells with an inclusion across the 154Q mice assessed. To determine whether the NLS mutation impacted the formation of nuclear inclusions in cerebral cortex, dorsal striatum, and CA1 of the hippocampus earlier in disease progression, these regions were assessed at 5 weeks of age (Figure 6I-K and Figure S6). At this age, there was a significant difference in in number of cells with a nuclear inclusion between *Atxn1(175Q)* and *Atxn1(175Q)K772T* mice for all three brain regions assessed. These findings indicate that the nuclear inclusion formation process is markedly delayed in mice with the NLS mutation. Additionally, the results show that the timing of nuclear inclusion formation is brain-region specific and not associated with the development of key disease phenotypes.

**Figure 6.**
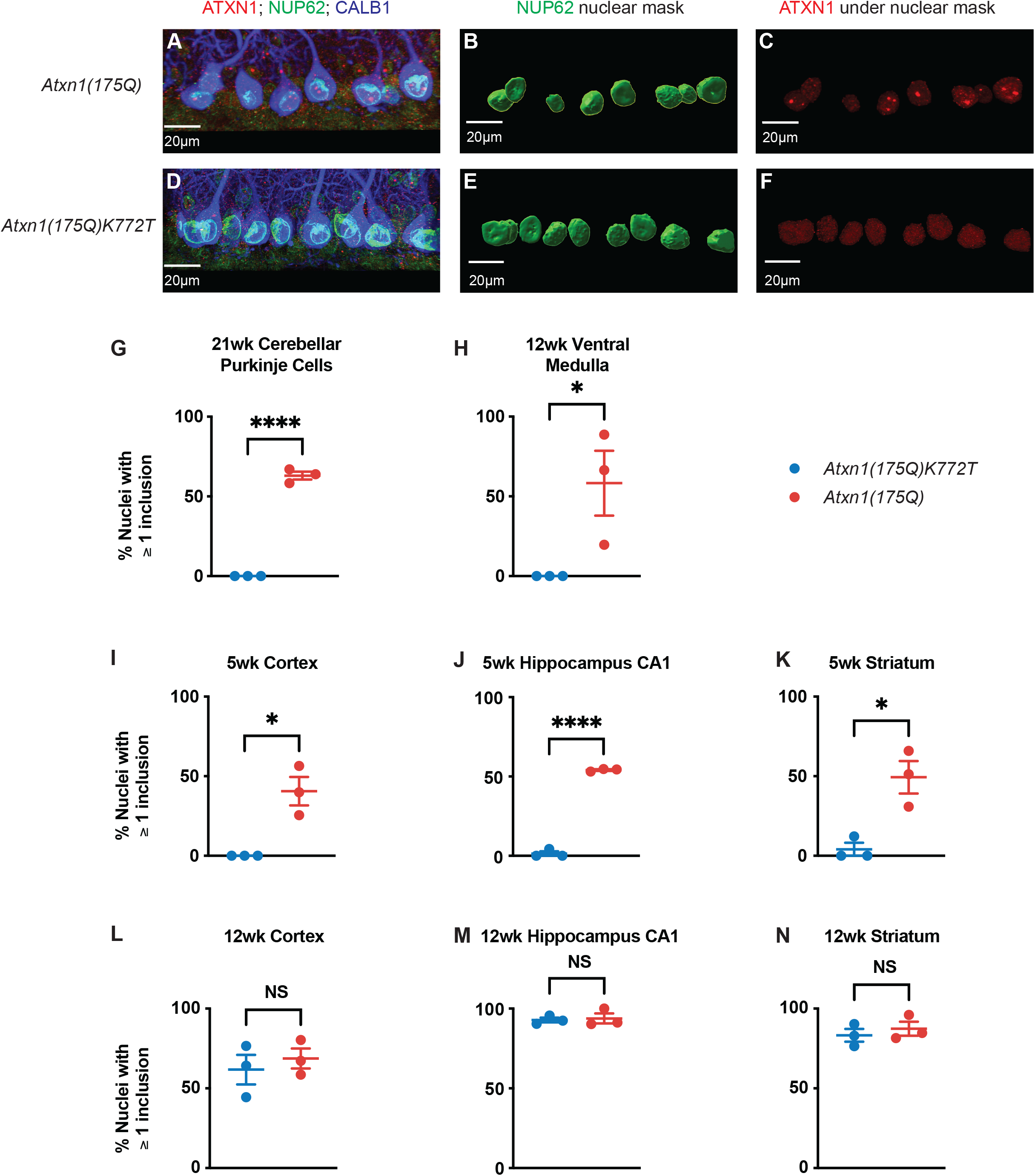
ATXN1 nuclear inclusions. (A and D) Immunofluorescent staining in cerebellar Purkinje cells of *Atxn1(175Q)* (A) and *Atxn1(175Q)K772T* (D) mice at 21 weeks of age. ATXN1 is shown in red, NUP62 is shown in green, and CALB1 is shown in blue. (B and E) Nuclear mask generated by Imaris using NUP62 staining in Purkinje cells of *Atxn1(175Q)* (B) and *Atxn1(175Q)K772T* (E) mice at 21 weeks of age. (C and F) ATXN1 staining under the nuclear mask in Purkinje cells of *Atxn1(175Q)* (C) and *Atxn1(175Q)K772T* (F) mice at 21 weeks of age. (G-N) Percentage of nuclei analyzed with at least one ATXN1 inclusion present in *Atxn1(175Q)* and *Atxn1(175Q)K772T* cerebellar Purkinje cells at 21 weeks (G), ventral medulla at 12 weeks (H), cerebral cortex at 5 weeks (I), CA1 of the hippocampus at 5 weeks (J), striatum at 5 weeks (K), cerebral cortex at 12 weeks (L), CA1 of the hippocampus at 12 weeks (M), and striatum at 12 weeks (N). For all brain regions at all time points, n=3 mice per genotype. Data are represented as mean ± SEM. Unpaired two-tailed t tests were performed. Significant results are denoted as * (p<0.05), ** (p<0.01), *** (p<0.001), and **** (p<0.0001). Statistical analysis details can be found in Table S1. *See also Figure S6*

### Mutation of the ATXN1[175Q] NLS restores the disease-associated transcriptomic expression profiles of brain regions in a regionally unique manner

Previous RNAseq analyses using the *Atxn1^154Q/2Q^* mouse model demonstrated that the transcriptomic profile associated with disease progression differs greatly between the cerebellum and medulla (Friedrich et al., 2018). To expand understanding of this regional specificity and evaluate the effect of the NLS mutation on genetic expression, RNAseq was performed on lysates from cerebellum, medulla, cortex, striatum, and hippocampus tissue in WT, *Atxn1(175Q)*, and *Atxn1(175Q)K772T* mice at 26 weeks of age. The number of significantly differentially expressed genes (DEGs) in the pairwise comparison between WT and *Atxn1(175Q)* mice was much larger than the number of DEGs in the pairwise comparison between WT and *Atxn1(175Q)K772T* for all brain regions assessed (Figure 7A). Although the cortex, striatum, and hippocampus had the largest total number of DEGs for both genotype comparisons, the cerebellum and medulla had the largest proportion of genes corrected by the NLS mutation. A gene was considered to be corrected by the NLS mutation if it was significantly differentially expressed in the comparison between WT and *Atxn1(175Q)* mice and not significantly differentially expressed in the comparison between WT and *Atxn1(175Q)K772T* mice. These results show that the NLS mutation has a substantial influence on transcriptomic profile throughout the brain, but its strongest impact seems to be on the cerebellum and medulla.

**Figure 7.**
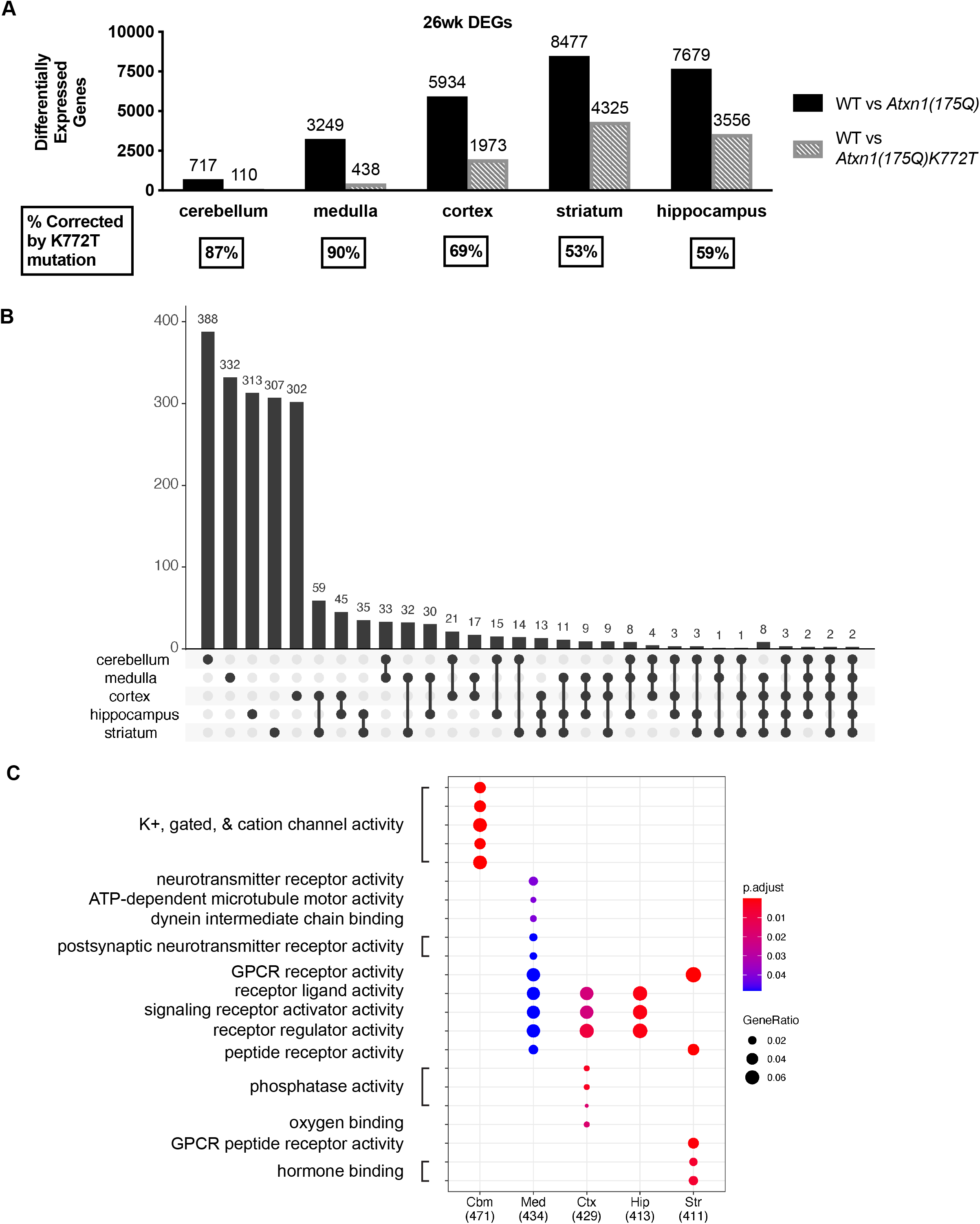
RNAseq analyses. (A) Significantly differentially expressed gene numbers between WT and *Atxn1(175Q)* mice (black) and between WT and *Atxn1(175Q)K772T* mice (gray) in cerebellum, medulla, cerebral cortex, striatum, and hippocampus tissue at 26 weeks of age. Genes that were significantly differentially expressed in the comparison between WT and *Atxn1(175Q)* mice and were not significantly differentially expressed in the comparison between WT and *Atxn1(175Q)K772T* mice were considered corrected by the K772T mutation. The percentage of genes corrected for each brain region is listed below the bar graph. For all genotypes, n=4 mice. (B) Brain region(s) where the top 500 corrected genes (genes with the largest absolute value Log_2_(Fold Change) in the comparison between WT and *Atxn1(175Q)* mice) from each region are found to be significantly differentially expressed. (C) Pathway enrichment analysis of the top 500 corrected genes from each brain region using the molecular function domain of the Gene Ontology database.

To further study the groups of corrected genes, we focused on the top 500 corrected genes in each brain region, defined as corrected genes with the largest absolute value Log_2_(Fold Change) in the comparison between WT and *Atxn1(175Q)* mice. The UpSetR package (Conway et al., 2017) was used to generate a diagram showing the number of top corrected genes from each brain region that were found to be unique to a particular region or shared by any combination of brain regions (Figure 7B). For all five brain regions assessed, the majority (>60%) of the top 500 corrected genes were unique to that brain region.

The top 500 corrected genes from each brain region were assessed for pathway enrichment using the clusterProfiler package (Yu et al., 2012) (Figure 7C). In looking specifically for enrichment of the pathways in the molecular function domain of the Gene Ontology database, the cerebellum stands out as having unique pathway enrichment with no overlap among the other brain regions assessed. While there is some overlap in pathway enrichment between the other brain regions, particularly between the cortex and hippocampus, most regions show a unique pathway enrichment. Together, these results support the idea that pathological mechanisms downstream of nuclear entry of expanded ATXN1 have regional specificity.

## DISCUSSION

While neurodegenerative diseases are often associated with prominent pathology in a certain brain region or neuronal population, many of these conditions impact diverse structures, circuitry, and cell populations that lack obvious pathology. In the case of SCA1, cerebellar Purkinje cell degeneration is a frequent and prominent pathological feature. Yet, SCA1 patients present with symptoms linked to multiple different brain regions (Koscik et al., 2020; Reetz et al., 2013). Previously, we showed that nuclear localization of expanded ATXN1 is critical for pathogenesis in cerebellar Purkinje cells (Klement et al., 1998). In this study, we assessed the importance of ATXN1 nuclear localization beyond the cerebellum and characterized its role in molecular aspects of disease in diverse brain regions associated with SCA1-like phenotypes. We found that mutating the NLS of expanded ATXN1[175Q] in a knock-in mouse model of SCA1 reduced nuclear localization of ATXN1[175Q] and dramatically mitigated a spectrum of phenotypes characteristic of SCA1.

### Severity of SCA1-like phenotypes are reduced in *ATXN1(175Q)K772T* mice

A complete rescue of rotarod performance was observed when the NLS mutation was introduced into ATXN1[82Q] expressed exclusively in Purkinje cells of a transgenic SCA1 mouse model (Klement et al., 1998). In contrast, while the rotarod performance of *Atxn1(175Q)K772T* mice was significantly improved compared to *Atxn1(175Q)* mice, there was not a complete rescue as *Atxn1(175Q)K772T* mice did not perform as well as WT mice at any age assessed. Notably, at 6 weeks of age, improvement of performance across successive trial days (i.e. motor learning) in naïve mice, was not restored in *Atxn1(175Q)K772T* mice (Figure 2A). This finding suggests that the inability to learn on the rotarod by *Atxn1(175Q)* mice is not solely due to Purkinje cell dysfunction induced by nuclear localization of ATXN1[175Q]. Conversely, the decline in motor performance with increasing age observed in *Atxn1(175Q)* mice was rescued in *Atxn1(175Q)K772T* mice (Figure 2B), indicating that nuclear localization of ATXN1[175Q] influences motor function over time. Whether this age-dependent loss in motor performance is due to nuclear localization of ATXN1[175Q] exclusively in Purkinje cells remains to be determined.

Mutation of the ATXN1[175Q] NLS nearly doubled the lifespan of *Atxn1(175Q)K772T* mice relative to *Atxn1(175Q)* mice (Figure 2C). This extension is, to our knowledge, the most substantial of any survival study results seen in animal models of SCA1 (Friedrich et al., 2018; Nitschke et al., 2021; Coffin et al., unpublished data). Thus, proper nuclear localization plays a critical role in the pathological dysfunction that underlies the premature lethality of *Atxn1(175Q)* mice. Similar to SCA1 patients, lethality in knock-in SCA1 mouse models is thought to be caused by complications due to bulbar/brain stem dysfunction leading to problems with breathing (Genis et al., 1995; Orengo et al., 2018; Sasaki et al., 1996). Intriguingly, the NLS mutation had a substantial impact on the time course of nuclear inclusion formation in the ventral brainstem of mice with expanded ATXN1 (Figure 6). Thus, we suggest that neurons of the brainstem in which dysfunction contributes to premature lethality in SCA1 are particularly responsive to the NLS mutation.

Across the other phenotypes assessed, *Atxn1(175Q)K772T* mice consistently performed significantly better than *Atxn1(175Q)* mice, but not as well as WT mice. This suggests that nuclear localization plays a critical role in a variety of different brain regions and wide range of disease-related metrics, but the NLS mutation alone does not completely halt pathogenesis. Subcellular fractionation results indicate that the NLS mutation significantly hindered but did not completely inhibit expanded ATXN1 from entering the nucleus (Figure 4). Thus, it is unclear at this time whether the observed partial restoration of WT phenotypes in Atxn1(175Q)K772T mice is due to reduction in nuclear expanded ATXN1 or aspects of pathogenesis occurring outside of the nucleus.

### ATXN1 extractability, subcellular localization, and nuclear inclusion formation in *ATXN1(175Q)K772T* mice

As noted previously, the extractability of expanded ATXN1 declined with age throughout the brain of *Atxn1(175Q)* mice (Figure 5 and Figure S4). The extractability of ATXN1[175Q]K772T also declined with age but was significantly greater than the extractability of ATXN1[175Q] in all brain regions and at all ages, suggesting that extractability of expanded ATXN1 decreases upon localization to the nucleus. Consistent with this concept is the strong correlation between the extraction of ATXN1[175Q]K772T and its cytoplasmic proportion at 26 weeks-of-age for all brain regions examined except the medulla (Figure 5G).

In the medulla, ATXN1[175Q]K772T extractability relative to its cytoplasmic proportion was much greater than in the other brain regions and was even greater than ATXN1[2Q]. We suggest that this finding likely reflects increased stability of the expanded protein relative to ATXN1[2Q] unique to the medulla. Regardless, analysis of the subcellular distribution of ATXN1[175Q]K772T indicates that the nuclear proportion in the medulla was very similar to the proportion of nuclear ATXN1[175Q]K772T in the other regions of the brain (Figure 4 and Figure S3). Moreover, at both 5 and 26 weeks of age, ATXN1[175Q] was more extractable from the medulla than the other regions of the brain (Figure 5 and Figure S4). This finding indicates that once in the nuclei of cells in the medulla, expanded ATXN1 remains more soluble than it does in the nuclei of cells in other brain regions.

Consistent with the subcellular fractionation data, the presence of ATXN1 nuclear inclusions with age in *Atxn1(175Q)K772T* mice supports the conclusion that the entry of mutant protein into the nucleus is not completely blocked by the NLS mutation. Although some expanded ATXN1 enters the nucleus in *Atxn1(175Q)K772T* mice, the age at which nuclear inclusions form is delayed relative to *Atxn1(175Q)* mice across all brain regions assessed (Figure 6). There is no detectable change in the nuclear proportion of ATXN1 in the cortex, hippocampus, medulla, or striatum of *Atxn1(175Q)K772T* mice between 5 and 40-42 weeks of age (Figure 4), but there is an increase in the proportion of cells with nuclear inclusions in these regions with age (Figure 6). Thus, we suggest that nuclear inclusions are a more sensitive marker of levels of expanded ATXN1 in the nucleus. The contribution of ATXN1 nuclear inclusions to the reduction in extractability of expanded ATXN1 remains unclear.

### Distinct brain region molecular disease-associated signatures

In a previous study, an antisense oligonucleotide targeting *Atxn1* RNA improved motor performance and survival of *Atxn1^154Q/2Q^* mice and RNAseq indicated that the molecular aspects of pathogenesis, as assessed by characterization of DEGs, differed between the cerebellum and medulla (Friedrich et al., 2018). Here, we found that proper nuclear localization of mutant ATXN1 is a seminal aspect of many SCA1-like CNS phenotypes including motor dysfunction, cognitive deficits, premature lethality, and brain weight. We utilized RNAseq to identify DEGs with expression corrected by the NLS K772T mutation in the cerebellum, medulla, hippocampus, cerebral cortex, and striatum. The NLS mutation substantially reduced the number of DEGs throughout the brain of *Atxn1(175Q)K772T* mice (Figure 6).

The largest proportional impact of the K772T NLS mutation on correcting DEGs was in the cerebellum (87% corrected) and medulla (90% corrected), which paralleled the strong age-related improvement of motor performance and extended survival associated with the NLS mutation. Moreover, the concept that disease-associated DEGs differ among brain regions in a SCA1 mouse model is supported by analyses of the top 500 genes corrected by the NLS mutation from each brain region. First, the vast majority of the top 500 corrected genes are unique to each brain region. Additionally, pathway enrichment analysis of the top 500 corrected genes from each brain reveled that, while there was some overlap in pathway enrichment between some brain regions, particularly between the cortex and hippocampus, most regions show a unique pathway enrichment profile. Notably, the cerebellar pathway was unique in its enrichment of genes associated with potassium, gated, and cation channel activity. This finding is consistent with recent work suggesting that early changes in expression of specific ionchannels and receptors are important for regulating membrane excitability and contribute to both motor dysfunction and structural changes in neurons that consistently precede cell death (Paulson et al., 2017). Overall, these results indicate that diseased-associated transcriptomic profiles are largely unique to each of the different brain regions assessed.

In summary, this study demonstrates that proper nuclear localization of expanded ATXN1 is an important aspect of pathogenesis across the spectrum of phenotypes characteristic of SCA1. Thus, it will be important to perform additional studies to better understand the regulation of ATXN1 nuclear localization. We argue that the results of this study indicate that inhibition of ATXN1 nuclear entry has considerable potential as a therapeutic approach for SCA1.

## ACKNOWLEDGEMENTS

This study was supported by NIH/NINDS grant RO1NS022920. The authors thank the Genomics Center and Mouse Behavior Core at the University of Minnesota.

## AUTHOR CONTRIBUTIONS

H.P.H. and H.T.O. conceived the study and wrote the paper. H.P.H., S.L.C., M.C, H.Y.Z, and H.T.O. designed experiments and interpreted the data. H.Y.Z. generated the *Atxn1^154Q/2Q^* mouse model. B.O. and H.P.H designed the CRISPR-Cas9 strategy used to generate the *Atxn1^175QK772T/2Q^* mouse model. B.R. assessed the efficacy of the CRISPR-Cas9 strategy in 3T3 cells. Y.Y. performed embryo injections and implantations. O.R. and S.S. bred mice and managed the colony. H.P.H. genotyped the mice. H.P.H., O.R., and S.S. conducted the survival study and animal weight measurements. O.R. and T.N.-M. performed behavioral assays. H.P.H. and B.O. performed subcellular fractionation assays. H.P.H., A.S., and M.R. performed extractability assays. L.D. sectioned and stained tissue for immunofluorescence. J.M. developed and implemented immunofluorescence image analyses. H.P.H. performed statistical analyses. H.P.H. and C.H. performed RNAseq analyses. All authors reviewed the manuscript and provided input.

## METHODS

### RESOURCE AVAILABILITY

#### Lead Contact

Further information and requests for resources and reagents should be directed to the lead contact, Harry T. Orr.

#### Materials Availability

The *Atxn1^175QK772T/2Q^* mouse line generated in this study has been deposited to xxxxxx.

#### Data and Code Availability

All RNA sequencing FASTQ data files have been deposited to SRA accession #xxxxxx. RNA sequencing counts data and differential expression data have been deposited to GEO accession #xxxxxx. Accession numbers are listed in the key resources table. This paper does not report original code. Any additional information required to reanalyze the data reported in this paper is available from the lead contact upon request.

### EXPERIMENTAL MODEL AND SUBJECT DETAILS

#### Mice

The University of Minnesota Institutional Animal Care and Use Committee approved all animal use protocols. All mice were housed and managed by Research Animal Resources under specific pathogen-free conditions in an Association for Assessment and Accreditation of Laboratory Animal Care International approved facility. The mice had unrestricted access to food and water except during behavioral testing. In all experiments, equal numbers of male and female mice were used. All mice were age matched within experiments and littermate controls were used when possible.

All mice were maintained on a C57BL/6 genetic background. WT *(Atxn1^2Q/2Q^)* mice were ordered from The Jackson Laboratory. *Atxn1^175Q/2Q^* mice are the result of spontaneous trinucleotide repeat expansion in the *Atxn1* gene of *Atxn1^154Q/2Q^* mice that were previously described (Watase et al., 2002).

#### Generation of *Atxn1^175QK772T/2Q^* mouse model

*Atxn1^175QK772T/2Q^* mice were generated via CRISPR-Cas9-mediated gene editing (Richardson et al., 2016; Yang et al., 2014). The guide RNA sequence (5’-CGACCACCTCCTCTTCCTCG-3’) was selected based on the highest on-target potential and the lowest off target risk using the Custom Alt-R CRISPR-Cas9 guide RNA design software (Integrated DNA Technologies). Based on guidance from the University of Minnesota Genome Engineering Shared Resource, this guide RNA (sgRNA) with a modified EZ scaffold was purchased from Synthego.

A single-stranded oligonucleotide (ssODN) was purchased from Integrated DNA Technologies as a template for homologous-directed recombination to introduce the K772T alteration and another synonymous mutation to endogenous *Atxn1* (5’-AAAATAGGATTGCCTGCAGCACCCTTCCTCAGCAAAATAGAACCGAGCAAACCCACAGCtACGAGGAccAGGAGGTGGTCGGCGCCGGAGACCCGTAAACTGGAGAAGTCGGAGGACGAGCCACCTTTGA-3’; modified nucleotides are lowercase). Two nucleotide modifications were required to introduce the K772T alteration. The other nucleotide alteration serves two functions: ablation of the PAM site and introduction of an Alu1 enzyme digest site used for genotyping.

Prior to injecting mouse embryos, accurate targeting of the sgRNA and introduction of the desired nucleotide alterations via homologous-directed recombination by the ssODN was confirmed by the University of Minnesota Genome Engineering Shared Resource in NIH3T3 cells.

At the University of Minnesota Mouse Genetics Laboratory (MGL), female C57BL/6 mice were super-ovulated via injection of pregnant mares serum gonadotropin followed by injection of human chorionic gonadotropin 48 hours later. The females were then bred to *Atxn1^175Q/2Q^* males overnight and zygotes were collected the following day. An injection mixture containing 39.6ng/μL Cas9 protein (Integrated DNA Technologies), 7ng/μL sgRNA, and 20ng/μL ssODN was prepared per MGL protocols and injected into the pronucleus of extracted zygotes. Zygotes were then transferred by MGL into oviducts of pseudopregnant females. Offspring resulting from this procedure were genotyped using the protocols in the following section. Integration of desired nucleotide changes on the expanded *Atxn1* allele was confirmed by Sanger sequencing. The mice were backcrossed for a minimum of 3 generations before conducting molecular or behavioral assays to avoid off-target effects.

#### Genotyping

PCR with the following primers was used to determine which animals have an expanded *Atxn1* allele: *Atxn1* 175Q Forward (5’-ACCTTCCAGTTCATTGGGTC-3’) and *Atxn1* 175Q Reverse (5’-GCTCTGTGGAGAGCTGGA-3’). To assess the presence of the K772T alteration, PCR was performed using the following primers: K772T Forward (5’-GCCGTGTTCCAAACTCTCTG-3’) and K772T Reverse (5’-GGTCTCTACTTGCCCACGTTA-3’) followed by an Alu1 restriction enzyme digest.

### METHOD DETAILS

#### Behavioral Analyses

Investigators were blinded to the genotype of the mice for all behavioral analyses. *Atxn1^2Q/2Q^, Atxn1^175Q/2Q^*, and *Atxn1^175QK772T/2Q^* mice were used in all behavioral tests. Mice were habituated to the testing room for at least 15 min prior to the start of testing on each day. All testing apparatus was cleaned between each animal with 70% ethanol.

##### Rotarod

The same cohort of mice was tested at 6, 12, 18, and 24 weeks of age. Mice were assessed on rotarod apparatus (Ugo Basile) using an accelerating protocol: 5 to 50 rpm, 5-min ramp duration, 5-min maximum trial length. The test consisted of a total of 4 trials per day for 4 consecutive days. Animals were segregated by sex during testing and run in consistent groups (up to 5 at a time). To ensure enough recovery time between trials, animals were given 10-15 min to rest between successive trials, which included the time used to test the other groups in the trial. A trial ended when an animal failed to stay on the rotarod or if they made 2 consecutive rotations clinging to the rod and not ambulating.

##### Barnes Maze

The same cohort of mice was assessed at 7 and 17 weeks of age. The maze is a circular table 92cm above the floor with a diameter of 92cm. The table has twenty circular holes located at equal distances around the perimeter, each with a diameter of 5 cm. One hole (the target hole) leads to a 5.7cm wide × 11.5cm long × 6.4cm deep escape chamber in which the animal can hide and the other 19 holes were closed. Illumination at the center of the table is maintained at 250 lux until the mouse enters the escape chamber to motivate it to search for the target hole. The testing room had visual cues on the walls to serve as landmarks. The position of each mouse was tracked using ANY-maze (Stoelting). Mice were exposed to the maze for 4 3-min trials per day (intertrial interval of approximately 10 min) during 4 consecutive training days. Mice that did not enter the escape box within 3 min were gently guided to it. To eliminate odor tracking and ensure mice are only using spatial cues, the maze was rotated 90° between each trial while maintaining the location of the escape box in relation to the rest of the room. Video footage of mice traveling to the escape chamber during training was analyzed using the Barnes-maze unbiased strategy (BUNS) algorithm (Illouz et al., 2016) to classify search strategies and score cognitive performance. Following training, a probe test was conducted with closed holes and no escape chamber. Each mouse was allowed to explore the maze freely for 90s. Latency to reach the blocked target hole, total distance travelled, speed, and time spent in each region of the maze were automatically recorded.

##### Contextual Fear Conditioning

Mice were assessed at 8 weeks of age. They were trained in a fear-conditioning chamber (Med Associates) that can deliver an electric shock paired with a tone. This device was located inside a soundproof box that contained a digital camera. Each mouse was placed individually in the chamber for habituation and left undisturbed for 3 min. Each mouse was given 5 fear acquisition stimuli foot-shocks (2s, 0.7mA) at varying intervals over a 7 min period (the shock timing pattern was consistent across all mice tested). The mouse was then returned to its home cage overnight. Contextual fear memory was assessed 24 hours after training. Mice were placed in the fear acquisition environment for 3 min without a foot-shock. To assess fear in a novel environment, mice were placed in the chamber, which had been modified (wall pattern, flooring, and scent) to distinguish it from the original context. For all training sessions and trials, mouse movement was recorded and analyzed using Video Freeze software (Med Associates). Freezing was scored only if the animal was immobile for at least 500ms.

#### RT-qPCR

Total RNA was isolated from cerebellum, medulla, cerebral cortex, hippocampus, and striatum tissue from *Atxn1^2Q/2Q^, Atxn1^175QK772T/2Q^* and *Atxn1^175Q/2Q^* mice by homogenizing it in 500μL TRIzol reagent (Thermo Fisher Scientific) per the manufacturer’s instructions. Synthesis of cDNA was performed in duplicate from 500ng RNA in 10μL reactions using the iScript Advanced cDNA Synthesis kit (Bio-Rad). Reactions were diluted 1:5 in water. qPCR was performed using 2μL cDNA in 10μL Probes Master (Roche) reactions on a Roche 480 Lightcycler. Target gene and reference gene reactions were amplified in separate wells under cycling conditions of 95°C for 10s, followed by 60°C for 10s for 35 cycles. Gapd was used as a reference gene. A complete list of primers and probes used for qPCR is presented in Table S2. Quantitation cycle (C_q_) values were determined using the Roche second derivative maximum calculation. Relative quantification was done using standard 2^Cq^.

#### Western Blots

##### Subcellular Fractionation

Cerebellum, medulla, cerebral cortex, hippocampus, and striatum tissue was collected from *Atxn1^175QK772T/2Q^* and *Atxn1^175Q/2Q^* mice at 5 weeks of age and from *Atxn1^175QK772T/2Q^* mice at 40-42 weeks of age. Cytoplasmic and nuclear protein extracts were collected using the NE-PER kit (Thermo Fisher Scientific). Protein concentrations were measured using the Pierce BCA Protein Assay kit (Thermo Fisher Scientific). Cytoplasmic and nuclear samples from each brain region containing 5μg total protein was boiled in Laemmli loading buffer and run on a 4%–20% Bio-Rad precast gel. Protein was transferred to a nitrocellulose membrane using the BioRad Trans-Blot Turbo system. Blots were cut at approximately 75kDa and blocked overnight at 4°C in 5% milk PBST (phosphate-buffered saline, 0.1% Tween 20). Blot sections were probed overnight at 4°C 1:5000 with the ATXN1 antibody 11750 (Servadio et al., 1995) or 1:2500 with H1 antibody (Invitrogen) diluted in 5% milk PBST. Blots were washed 3 times with PBST and then placed in 5% milk PBST plus 1:2500 rabbit specific horseradish peroxidase (HRP) antibodies (GE Healthcare) at room temperature for 4 hours. Blots were washed 3 times with PBST followed by Super Signal West Dura (Thermo Fisher Scientific) detection reagents and imaged on an ImageQuant LAS 4000. The H1 blots were stripped with Restore Western Blot Stripping Buffer (Thermo Fisher Scientific) for 15 min at room temperature, washed 3 times with PBST, and blocked overnight at 4°C in 5% milk PBST. These blots were then probed overnight at 4°C 1:1000 with GAPDH antibody (EMD Millipore Corp), washed 3 times with PBST, placed in 5% milk PBST plus 1:2500 mouse specific HRP antibodies (GE Healthcare) at room temperature for 4 hours, and visualized via the same method as the ATXN1 and H1 blots.

##### ATXN1 Extractability

Cerebellum, medulla, cerebral cortex, hippocampus, and striatum tissue was collected from *Atxn1^2Q/2Q^, Atxn1^175QK772T/2Q^* and *Atxn1^175Q/2Q^* mice at 5 weeks and 26 weeks of age. Samples were homogenized using a tissue grinder in an appropriate volume (350μl for medulla and striatum, 500μl for cerebellum, cortex, and hippocampus) of Tris Triton lysis buffer (50mM Tris, pH 7.5, 100mM NaCl, 2.5mM MgCl2, 0.5% Triton X-100) that included MilliporeSigma protease inhibitors II and III and a Roche Complete Mini Protease inhibitor tablet. Homogenized samples were shaken at 1500rpm at 4°C for 1 hour, frozen and thawed in liquid nitrogen and 37°C water bath 3 times, and centrifuged at 21,000xg for 10 min at 4°C. Samples containing 30μg total protein were boiled in Laemmli loading buffer and run on a 4%–20% Bio-Rad precast gel. Protein was transferred to a nitrocellulose membrane using the BioRad Trans-Blot Turbo system. Blots were cut at approximately 75kDa and blocked overnight at 4°C in 5% milk PBST (phosphate-buffered saline, 0.1% Tween 20). Blots were probed overnight at 4°C 1:5000 with the ATXN1 antibody 11750 (Servadio et al., 1995) or 1:10,000 with α-Tubulin antibody (MilliporeSigma) diluted in 5% milk PBST. Blots were washed 3 times with PBST. ATXN1 blots were then placed in 5% milk PBST plus 1:2500 rabbit specific HRP antibodies (GE Healthcare) while α-Tubulin blots were placed in 5% milk PBST plus 1:10,000 mouse specific HRP antibodies (GE Healthcare) at room temperature for 4 hours. Blots were washed 3 times with PBST and then ATXN1 blots were washed with Super Signal West Dura (Thermo Fisher Scientific) while α-Tubulin blots were washed with Super Signal West Pico (Thermo Fisher Scientific) detection reagents. Blots were imaged on an ImageQuant LAS 4000.

#### Immunofluorescence

Mice were deeply anesthetized with Ketamine, transcardially exsanguinated with PBS, and perfused using 10% formalin. Brains were post-fixed overnight in 10% formalin then stored in PBS at 4°C until sectioning. Sagittal sections of 50μm were isolated using a Leica VT 1000S vibratome. Sections were permeabilized in 1% Triton X-100 in PBS. Sections were blocked for 1 hour in 5% normal donkey serum and 0.3% Triton X-100 in PBS. Subsequent staining was carried out in 2% normal donkey serum and 0.3% Triton X-100 in PBS. Sections were incubated for 24 hours with primary antibodies (1:1000 anti-mouse NUP62 (BD Biosciences), 1:500 anti-rabbit 12NQ (Pérez Ortiz et al., 2018), 1:250 anti-guinea pig CALB1 (Synaptic Systems)) at 4°C. Following incubation, sections were washed four times in PBS and exposed to secondary antibodies (Jackson Immunoresearch Labs Alexa Fluor 488 anti-mouse, Cy3 anti-rabbit, and Alexa Fluor 647 anti-guinea pig) for 24 hours at 4°C. Sections were washed four times in PBS and mounted using ProLong Gold antifade reagent (Thermo Fisher Scientific). Fluorescently labeled tissue was imaged with 488/559/635nm lasers using an Olympus Fluoview 1000 IX2 inverted confocal microscope with a 40x oil immersion objective (NA 1.3) with an xy voxel size of 0.305-0.450μm and z-step size of 0.84μm. Fluorescent emissions were detected with photomultiplier tubes with spectral ranges of 500-545nm (Alexa Fluor 488), 575-620nm (Cy3), and 655-755nm (Alexa Fluro 647) with the confocal aperture set to 85μm. Images from each brain region were captured using identical acquisition settings for *Atxn1^175QK772T/2Q^* and *Atxn1^175Q/2Q^* samples. Nuclear ATXN1 inclusions were analyzed using batched processes developed with Imaris (Oxford Instruments, v9.8). The Imaris surfaces module was used to isolate nuclear ATXN1 staining by creating a nuclear mask from NUP62 staining and to identify nuclear ATXN1 inclusions. Inclusion identification used identical thresholding parameters for *Atxn1^175QK772T/2Q^* and *Atxn1^175Q/2Q^* samples and was adjusted between tissue regions to reduce background detection. Inclusion identification was batched for all images from the same tissue region and background was further eliminated using a 1um^3^ volume filter. Incomplete or “chopped” inclusions were removed with a z-filter. Complete inclusions were analyzed by volume and mean fluorescent intensity of the filtered inclusion surfaces. Inclusion frequency was determined by using the Imaris spots module to count nuclear ATXN1 inclusions and NUP62-labled nuclei.

#### RNA Extraction and Sequencing

Cerebellum, medulla, cerebral cortex, hippocampus, and striatum tissue was isolated 26-week-old *Atxn1^2Q/2Q^, Atxn1^175QK772T/2Q^*, and *Atxn1^175Q/2Q^* mice and stored in *RNAlater* solution (Thermo Fisher Scientific). Total RNA was isolated using TRIzol reagent (Thermo Fisher Scientific) following the manufacturer’s protocols. Tissue was homogenized using RNase-Free disposable pellet pestles in a motorized chuck. Purified RNA was sent to the University of Minnesota Genomics Center for quality control, including quantification using fluorimetry via RiboGreen assay kit (Thermo Fisher Scientific) and RNA integrity was assessed via capillary electrophoresis using an Agilent BioAnalyzer 2100 to generate an RNA integrity number (RIN). RIN values for submitted RNA were above 8.0 for all samples except one medulla sample (RIN = 6.8). All submitted RNA samples had greater than 1μg total mass. Library creation was completed using oligo-dT purification of polyadenylated RNA, which was reverse transcribed to create cDNA. cDNA was fragmented, blunt ended, and ligated to barcode adaptors. Libraries were size selected to 320 bp ± 5% to produce average inserts of approximately 200 bp, and size distribution was validated using capillary electrophoresis and quantified using fluorimetry (PicoGreen, Thermo Fisher Scientific) and qPCR. Libraries were then normalized, pooled, and sequenced on an S4 flow cell by an Illumina NovaSeq 6000 using a 150-nucleotide, paired-end read strategy. The resulting FASTQ files were trimmed, aligned to the mouse reference genome (GRCm38), sorted, and counted using the Bulk RNAseq Analysis Pipeline from the Minnesota Supercomputing Institute’s Collection of Hierarchical UMII/RIS Pipelines (v0.2.0) (Baller et al., 2019). Genes less than 300 bp are too small to be accurately captured in standard RNAseq library preparations, so they were discarded from all downstream analyses.

#### RNAseq Analyses

Differential gene expression analysis was performed using the edgeR package (McCarthy et al., 2012; Robinson et al., 2009) (v3.30.3) in R (R Foundation for Statistical Computing v3.6.1). All five brain regions were analyzed independently. Genes with fewer than 10 counts across all samples in each region were excluded. Genes with FDR values less than or equal to 0.05 were considered significant. UpSet plots were created using the UpSetR package (Conway et al., 2017) (v1.4.0) in R (v4.0.3). Pathway analysis and dot plot creation was performed using the clusterProfiler package (Yu et al., 2012) (v3.16.1) in R (v4.0.3).

### QUANTIFICATION AND STATISTICAL ANALYSIS

For all experiments, pre-hoc power calculations were performed in MATLAB (MathWorks) to determine the number of mice per genotype necessary to reach 80% power with an alpha level of 0.05. Historical data comparing *Atxn1^2Q/2Q^* mice to *Atxn1^154Q/2Q^* mice was used to estimate effect size (Cohen’s d) for these power estimates. Aside from power calculations performed in MATLAB and the RNAseq analyses performed in R, all other statistical tests were performed using GraphPad Prism software (v9.0). All data were checked for normality and met assumptions for using parametric statistical tests. For multigroup comparisons, one-way or two-way ANOVAs were performed. When working with repeated-measures data, sphericity was not assumed; thus, a Geisser-Greenhouse correction was used. When ANOVA findings were significant, (p<0.05), the analysis was followed by multiple comparisons testing using Tukey’s (all pairwise comparisons) or Dunnett’s (comparisons back to one control group) correction for multiple comparisons. Survival analysis, plotted as Kaplan-Meyer curves, was assessed using log-rank Mantel-Cox and Gehan-Breslow-Wilcoxon tests. Western blots were quantified using ImageQuant TL software (Cytiva). For subcellular fractionation data, the nuclear proportion of expanded ATXN1 was calculated for each animal by dividing the intensity of the expanded ATXN1 nuclear band by the combined intensity of the nuclear band plus the cytoplasmic band. ATXN1 extractability was calculated by dividing the expanded ATXN1 band by the ATXN1[2Q] band for each animal. All data presented are shown as mean ± SEM. Significant results are denoted as * (p<0.05), ** (p<0.01), *** (p<0.001), and **** (p<0.0001). Complete statistical analysis details for all data analyzed in Prism are presented in Table S1.

## SUPPLEMENTAL FIGURES

**Figure S1. Cerebellar RT-qPCR expression of genes associated with disease progression in Purkinje cells, *Related to Figure 2***

(A-C) Relative expression of genes normalized to *Gapdh* in cerebellar Purkinje cells at 12 (A), 18 (B), and 26 (C) weeks of age. Data are represented as mean ± SEM. One-way ANOVAs with Dunnett’s post hoc test relative to WT expression were performed. Significant results are denoted as * (p<0.05), ** (p<0.01), *** (p<0.001), and **** (p<0.0001). Statistical analysis details can be found in Table S1.

*See also Table S2*

**Figure S2. Contextual fear conditioning, *Related to Figure 3***

Contextual fear conditioning assessment freezing time percentage among naïve mice at 8 weeks of age. Data are represented as mean ± SEM and n values for each genotype are shown in the bottom of each bar. A one-way ANOVA with Tukey’s post hoc test was performed.

Significant results are denoted as * (p<0.05), ** (p<0.01), *** (p<0.001), and **** (p<0.0001). Statistical analysis details can be found in Table S1.

**Figure S3. Subcellular fractionation Western blots, *Related to Figures 4 and 5G***

(A-H) Subcellular fractionation Western blot used to quantify nuclear proportion of expanded ATXN1 in *Atxn1(175Q)* and *Atxn1(175Q)K772T* mice at 5 weeks of age and in medulla (A), cortex (C), hippocampus (E), and striatum (G) and in *Atxn1(175Q)K772T* mice at 40-42 weeks of age in the medulla (B), cortex (D), hippocampus (F), and striatum (H). For all brain regions at 5 weeks of age (A, C, E, and G), n=4 mice per genotype. For 40 week medulla (B), n=6. For 42 week cortex (D), hippocampus (F), and striatum (H), n=4.

**Figure S4. ATXN1 extractability, *Related to Figure 5***

(A, D, and G) Cortex (A), hippocampus (D), and striatum (G) extractability of expanded ATXN1 protein products relative to ATXN1[2Q] in *Atxn1(175Q)* and *Atxn1(175Q)K772T* mice at 5 and 26 weeks of age. Extractability was determined for each animal by dividing intensity of the expanded ATXN1 band by the intensity of the ATXN1[2Q] band. Data are represented as mean ± SEM and n=3 mice per genotype at each age. Two-way repeated measures ANOVAs with Tukey’s post hoc test were performed. Significant results are denoted as * (p<0.05), ** (p<0.01), *** (p<0.001), and **** (p<0.0001). Statistical analysis details can be found in Table S1.

(B and C) Western blots used for extractability quantification of expanded ATXN1 protein products from cortex lysates in *Atxn1(175Q)* and *Atxn1(175Q)K772T* mice at 5 weeks of age (B) and 26 weeks of age (C).

(E and F) Western blots used for extractability quantification of expanded ATXN1 protein products from hippocampus lysates in *Atxn1(175Q)* and *Atxn1(175Q)K772T* mice at 5 weeks of age (E) and 26 weeks of age (F).

(H and I) Western blots used for extractability quantification of expanded ATXN1 protein products from striatum lysates in *Atxn1(175Q)* and *Atxn1(175Q)K772T* mice at 5 weeks of age (H) and 26 weeks of age (I).

**Figure S5. Expanded ATXN1 protein quantification from extractability and subcellular fractionation Western blots, *Related to Figures 4, 5, S3, and S4***

(A-J) Quantification of total expanded ATXN1 in *Atxn1(175Q)K772T* and *Atxn1(175Q)* mice at 5 weeks of age from extractability Western blots and subcellular fractionation Western blots respectively in the cerebellum (A-B), cerebral cortex (C-D), hippocampus (D-E), medulla (F-G), and striatum (H-I). For all brain regions, n=3 mice per genotype for extractability blots and n=4 mice per genotype for subcellular fractionation blots. Data are represented as mean ± SEM.

Unpaired two-tailed t tests were performed. Significant results are denoted as * (p<0.05), ** (p<0.01), *** (p<0.001), and **** (p<0.0001). Statistical analysis details can be found in Table S1.

**Figure S6. Nuclear ATXN1, *Related to Figure 6***

(A-H) Representative images showing ATXN1 expression in the nucleus of cells in ventral medulla at 12 weeks of age from *Atxn1(175Q)* mice (A) and *Atxn1(175Q)K772T* mice (B), cortex at 5 weeks of age from *Atxn1(175Q)* mice (C) and *Atxn1(175Q)K772T* mice (D), hippocampus CA1 at 5 weeks of age from *Atxn1(175Q)* mice (E) and *Atxn1(175Q)K772T* mice (F), and striatum at 5 weeks of age from *Atxn1(175Q)* mice (G) and *Atxn1(175Q)K772T* mice (H).

Identical image acquisition settings were used for both genotypes at 12 weeks of age (A-B) and identical image acquisition settings were used for all images at 5 weeks of age (B-H).

## SUPPLEMENTAL TABLES

**Table S1. Detailed statistical summary of results, *Related to Figures 2, 3, 4, 5, 6, S1, S2, and S4***

A statistical summary of all data analyzed in Prism throughout the text.

**Table S2. List of oligonucleotides and probes used for RT-qPCR in this study, *Related to Figure1 and S2***

Oligonucleotides used for RT-qPCR quantification of gene expression.

