## Supplemental data for "Disrupting ATXN1 Nuclear Localization in a Knock-in SCA1 Mouse Model Improves a Spectrum of SCA1-Like Phenotypes and their Brain Region Associated Transcriptomic Profiles"

### SUPPLEMENTAL FIGURES

**Figure S1. Cerebellar RT-qPCR expression of genes associated with disease progression in Purkinje cells, *Related to Figure 2***

(A-C) Relative expression of genes normalized to *Gapdh* in cerebellar Purkinje cells at 12 (A), 18 (B), and 26 (C) weeks of age. Data are represented as mean ± SEM. One-way ANOVAs with Dunnett's post hoc test relative to WT expression were performed. Significant results are denoted as * ($p<0.05$), ** ($p<0.01$), *** ($p<0.001$), and **** ($p<0.0001$). Statistical analysis details can be found in Table S1.

*See also Table S2*

**Figure S2. Contextual fear conditioning, *Related to Figure 3***

Contextual fear conditioning assessment freezing time percentage among naïve mice at 8 weeks of age. Data are represented as mean ± SEM and n values for each genotype are shown in the bottom of each bar. A one-way ANOVA with Tukey's post hoc test was performed. Significant results are denoted as * ($p<0.05$), ** ($p<0.01$), *** ($p<0.001$), and **** ($p<0.0001$). Statistical analysis details can be found in Table S1.

**Figure S3. Subcellular fractionation Western blots, *Related to Figures 4 and 5G***

(A-H) Subcellular fractionation Western blot used to quantify nuclear proportion of expanded ATXN1 in *Atxn1(175Q)* and *Atxn1(175Q)K772T* mice at 5 weeks of age and in medulla (A), cortex (C), hippocampus (E), and striatum (G) and in *Atxn1(175Q)K772T* mice at 40-42 weeks of age in the medulla (B), cortex (D), hippocampus (F), and striatum (H). For all brain regions at 5 weeks of age (A, C, E, and G), n=4 mice per genotype. For 40 week medulla (B), n=6. For 42 week cortex (D), hippocampus (F), and striatum (H), n=4.

**Figure S4. ATXN1 extractability, *Related to Figure 5***

(A, D, and G) Cortex (A), hippocampus (D), and striatum (G) extractability of expanded ATXN1 protein products relative to ATXN1[2Q] in *Atxn1(175Q)* and *Atxn1(175Q)K772T* mice at 5 and 26 weeks of age. Extractability was determined for each animal by dividing intensity of the expanded ATXN1 band by the intensity of the ATXN1[2Q] band. Data are represented as mean ± SEM and n=3 mice per genotype at each age. Two-way repeated measures ANOVAs with Tukey's post hoc test were performed. Significant results are denoted as * (p<0.05), ** (p<0.01), *** (p<0.001), and **** (p<0.0001). Statistical analysis details can be found in Table S1.

(B and C) Western blots used for extractability quantification of expanded ATXN1 protein products from cortex lysates in *Atxn1(175Q)* and *Atxn1(175Q)K772T* mice at 5 weeks of age (B) and 26 weeks of age (C).

(E and F) Western blots used for extractability quantification of expanded ATXN1 protein products from hippocampus lysates in *Atxn1(175Q)* and *Atxn1(175Q)K772T* mice at 5 weeks of age (E) and 26 weeks of age (F).

(H and I) Western blots used for extractability quantification of expanded ATXN1 protein products from striatum lysates in *Atxn1(175Q)* and *Atxn1(175Q)K772T* mice at 5 weeks of age (H) and 26 weeks of age (I).

**Figure S5. Expanded ATXN1 protein quantification from extractability and subcellular fractionation Western blots, *Related to Figures 4, 5, S3, and S4***

(A-J) Quantification of total expanded ATXN1 in *Atxn1(175Q)K772T* and *Atxn1(175Q)* mice at 5 weeks of age from extractability Western blots and subcellular fractionation Western blots respectively in the cerebellum (A-B), cerebral cortex (C-D), hippocampus (D-E), medulla (F-G), and striatum (H-I). For all brain regions, n=3 mice per genotype for extractability blots and n=4 mice per genotype for subcellular fractionation blots. Data are represented as mean ± SEM.

Unpaired two-tailed t tests were performed. Significant results are denoted as * ($p<0.05$), ** ($p<0.01$), *** ($p<0.001$), and **** ($p<0.0001$). Statistical analysis details can be found in Table S1.

**Figure S6. Nuclear ATXN1, *Related to Figure 6***

(A-H) Representative images showing ATXN1 expression in the nucleus of cells in ventral medulla at 12 weeks of age from *Atxn1(175Q)* mice (A) and *Atxn1(175Q)K772T* mice (B), cortex at 5 weeks of age from *Atxn1(175Q)* mice (C) and *Atxn1(175Q)K772T* mice (D), hippocampus CA1 at 5 weeks of age from *Atxn1(175Q)* mice (E) and *Atxn1(175Q)K772T* mice (F), and striatum at 5 weeks of age from *Atxn1(175Q)* mice (G) and *Atxn1(175Q)K772T* mice (H). Identical image acquisition settings were used for both genotypes at 12 weeks of age (A-B) and identical image acquisition settings were used for all images at 5 weeks of age (B-H).

### SUPPLEMENTAL TABLES

**Table S1. Detailed statistical summary of results, *Related to Figures 2, 3, 4, 5, 6, S1, S2, and S4***

A statistical summary of all data analyzed in Prism throughout the text.

**Table S2. List of oligonucleotides and probes used for RT-qPCR in this study, *Related to Figure1 and S2***

Oligonucleotides used for RT-qPCR quantification of gene expression.

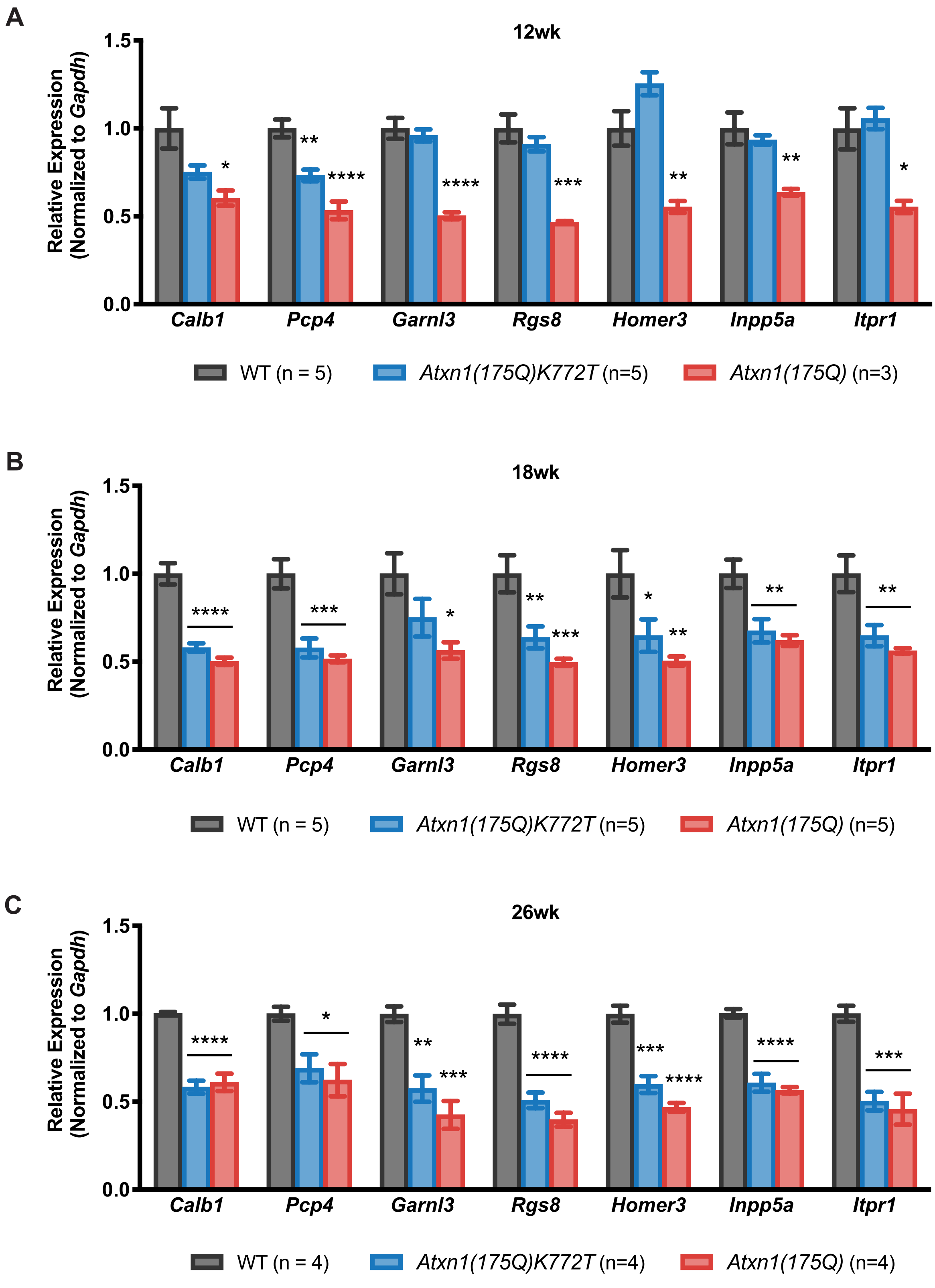


**Figure S1. Cerebellar RT-qPCR expression of genes associated with disease progression in Purkinje Cells.**

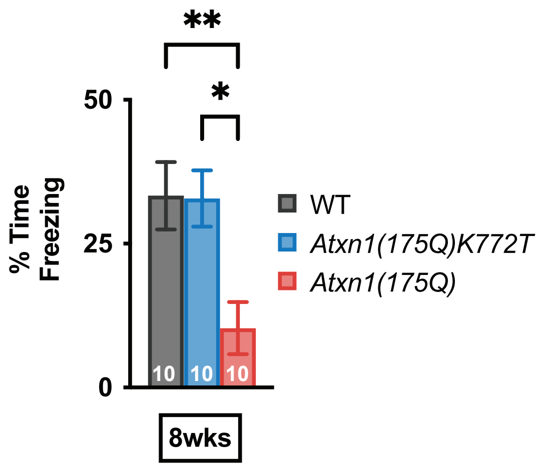


**Figure S2. Contextual fear conditioning**

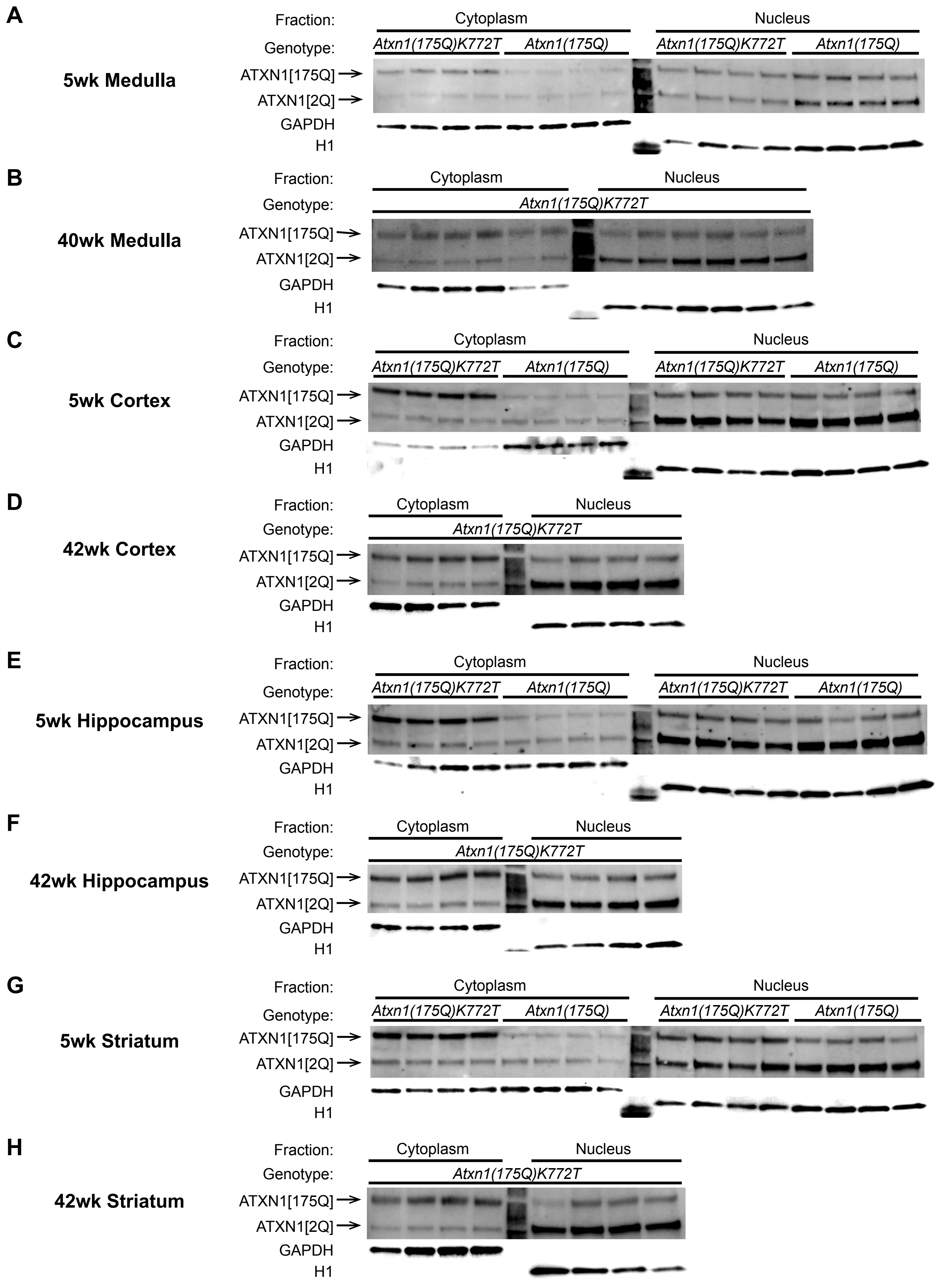


**Figure S3. Subcellular fractionation Western blots**

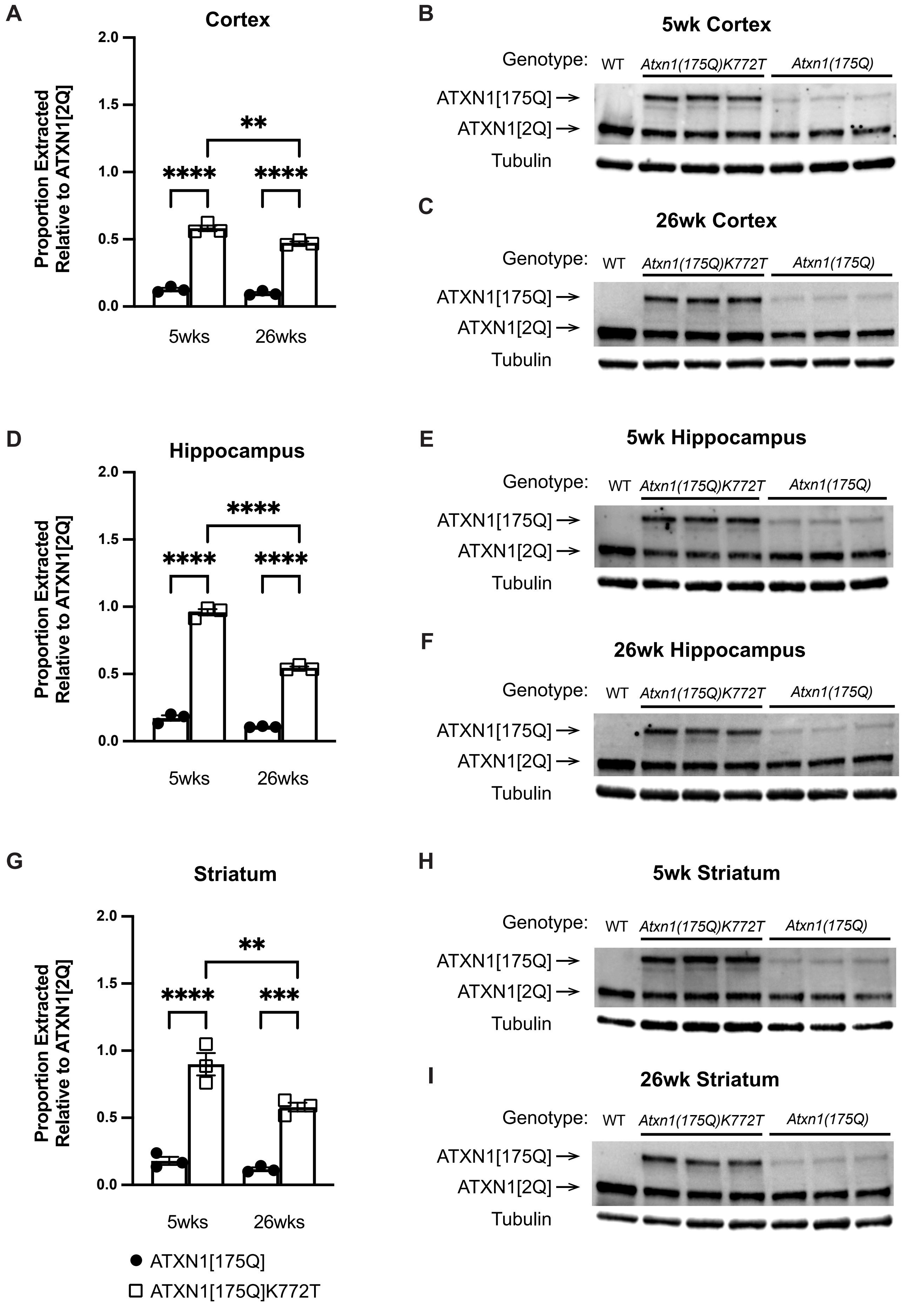


**Figure S4. ATXN1 extractability**

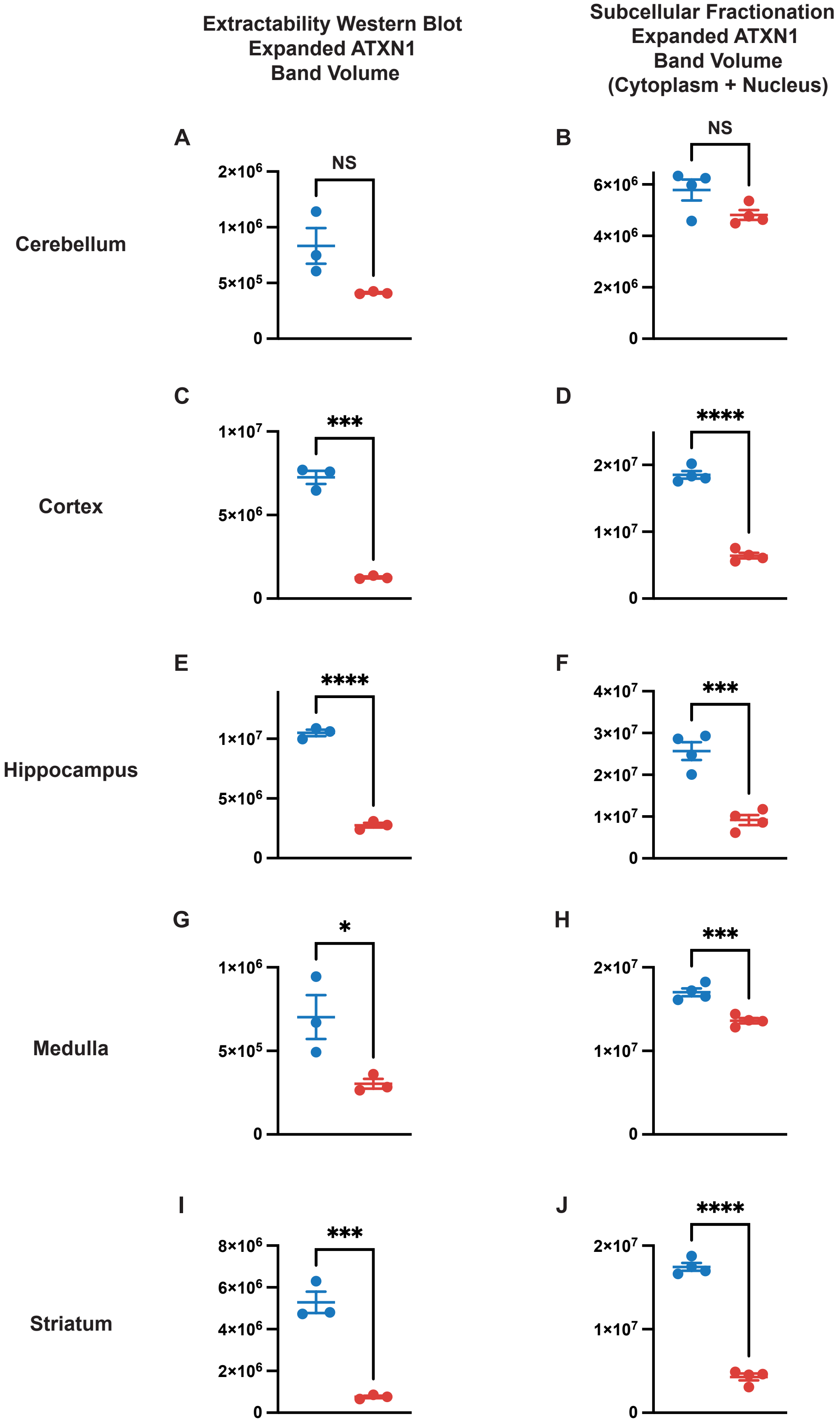


**Figure S5. Expanded ATXN1 protein quantification from extractability and subcellular fractionation Western blots**

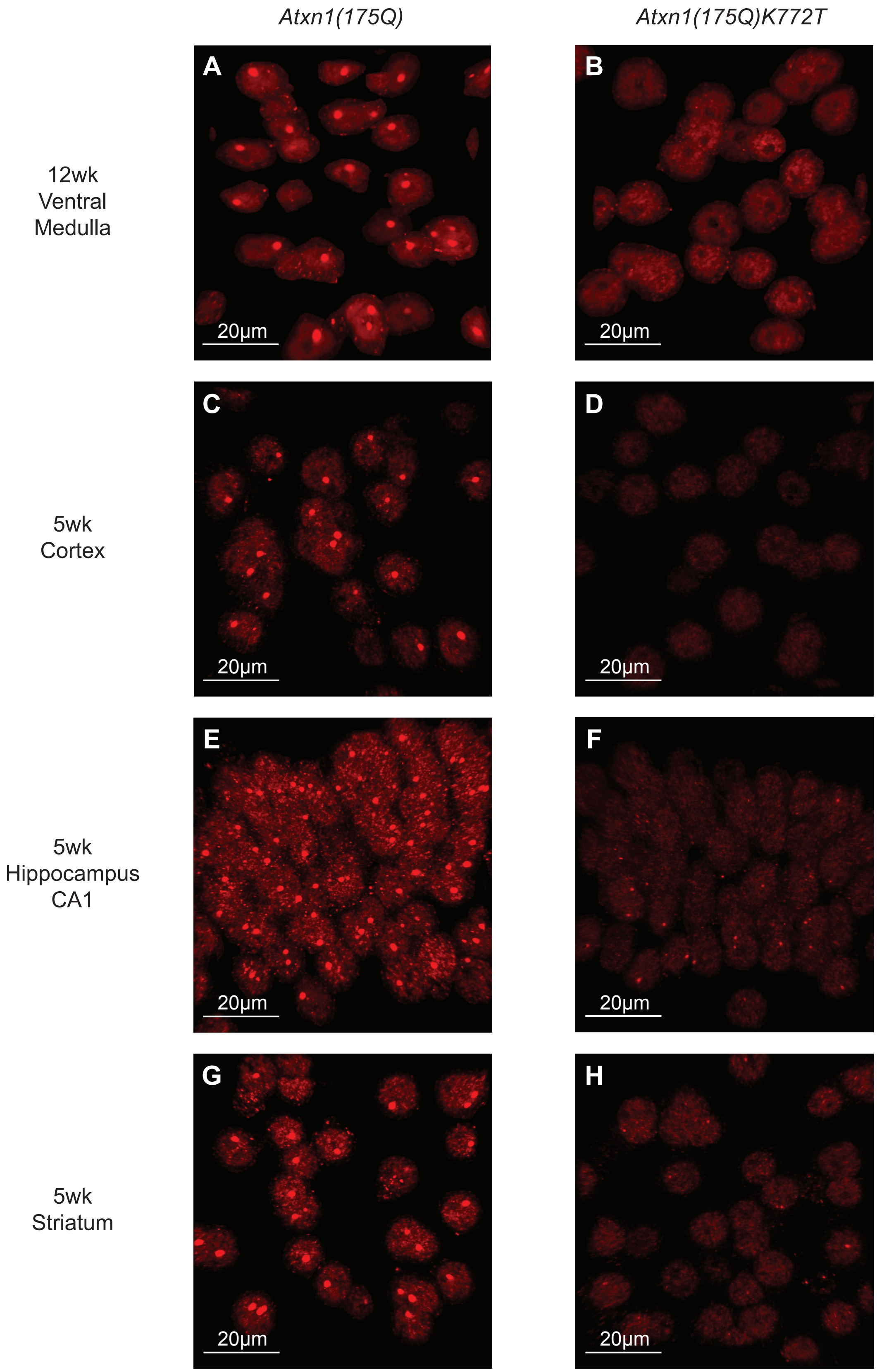


**Figure S6. Nuclear ATXN1**

| Figure | Assay | Measurement | Statistical Test | N(n) | Comparison | F (DFn, DFd) | *P* value |
|---|---|---|---|---|---|---|---|
| Fig. 1C | qPCR - 26wks | Atxn1 relative expression | Two-way RM ANOVA | 4,4,4 | Brain Region | F (2.113, 19.02) = 27.43 | <0.0001 |
| | | | | | Genotype | F (2, 9) = 5.442 | 0.0282 |
| | Cerebellum | | Dunnett's multiple comparisons test | | $Atxn1^{2Q/2Q}$ vs. $Atxn1^{175Q/2Q}$ | | 0.0585 |
| | | | | | $Atxn1^{2Q/2Q}$ vs. $Atxn1^{175QK772T/2Q}$ | | 0.4424 |
| | Medulla | | | | $Atxn1^{2Q/2Q}$ vs. $Atxn1^{175Q/2Q}$ | | 0.4965 |
| | | | | | $Atxn1^{2Q/2Q}$ vs. $Atxn1^{175QK772T/2Q}$ | | 0.1877 |
| | Cortex | | | | $Atxn1^{2Q/2Q}$ vs. $Atxn1^{175Q/2Q}$ | | 0.2736 |
| | | | | | $Atxn1^{2Q/2Q}$ vs. $Atxn1^{175QK772T/2Q}$ | | 0.3452 |
| | Hippocampus | | | | $Atxn1^{2Q/2Q}$ vs. $Atxn1^{175Q/2Q}$ | | 0.5143 |
| | | | | | $Atxn1^{2Q/2Q}$ vs. $Atxn1^{175QK772T/2Q}$ | | 0.1020 |
| | Striatum | | | | $Atxn1^{2Q/2Q}$ vs. $Atxn1^{175Q/2Q}$ | | 0.0489 |
| | | | | | $Atxn1^{2Q/2Q}$ vs. $Atxn1^{175QK772T/2Q}$ | | 0.0490 |
| Fig. 2A | Rotarod - 6wks | Latency to fall | Two-way RM ANOVA | 12,12,11 | Genotype | F (2, 32) = 8.942 | 0.0008 |
| | Trial 1 | | Tukey's multiple comparisons test | | $Atxn1^{2Q/2Q}$ vs. $Atxn1^{175Q/2Q}$ | | 0.7120 |
| | | | | | $Atxn1^{2Q/2Q}$ vs. $Atxn1^{175QK772T/2Q}$ | | 0.7560 |
| | | | | | $Atxn1^{175Q/2Q}$ vs. $Atxn1^{175QK772T/2Q}$ | | 0.2956 |
| | Trial 2 | | | | $Atxn1^{2Q/2Q}$ vs. $Atxn1^{175Q/2Q}$ | | 0.0778 |
| | | | | | $Atxn1^{2Q/2Q}$ vs. $Atxn1^{175QK772T/2Q}$ | | 0.0405 |
| | | | | | $Atxn1^{175Q/2Q}$ vs. $Atxn1^{175QK772T/2Q}$ | | 0.9565 |
| | Trial 3 | | | | $Atxn1^{2Q/2Q}$ vs. $Atxn1^{175Q/2Q}$ | | 0.0161 |
| | | | | | $Atxn1^{2Q/2Q}$ vs. $Atxn1^{175QK772T/2Q}$ | | 0.0021 |
| | | | | | $Atxn1^{175Q/2Q}$ vs. $Atxn1^{175QK772T/2Q}$ | | 0.9783 |
| | Trial 4 | | | | $Atxn1^{2Q/2Q}$ vs. $Atxn1^{175Q/2Q}$ | | <0.0001 |
| | | | | | $Atxn1^{2Q/2Q}$ vs. $Atxn1^{175QK772T/2Q}$ | | 0.0002 |
| | | | | | $Atxn1^{175Q/2Q}$ vs. $Atxn1^{175QK772T/2Q}$ | | 0.8777 |
| | Rotarod - 6wks | Latency to fall | Two-way RM ANOVA | 12,12,11 | Trial Day | F (2.628, 84.09) = 6.766 | 0.0007 |
| | $Atxn1^{2Q/2Q}$ | | Tukey's multiple comparisons test | | 1 vs 2 | | 0.0002 |
| | | | | | 1 vs 3 | | 0.0017 |
| | | | | | 1 vs 4 | | 0.0008 |
| | | | | | 2 vs 3 | | 0.1435 |
| | | | | | 2 vs 4 | | 0.1650 |
| | | | | | 3 vs 4 | | 0.9998 |
| | $Atxn1^{175QK772T/2Q}$ | | | | 1 vs 2 | | >0.9999 |
| | | | | | 1 vs 3 | | 0.9443 |
| | | | | | 1 vs 4 | | 0.0720 |
| | | | | | 2 vs 3 | | 0.9379 |
| | | | | | 2 vs 4 | | 0.1125 |
| | | | | | 3 vs 4 | | 0.1023 |
| | $Atxn1^{175Q/2Q}$ | | | | 1 vs 2 | | 0.6710 |
| | | | | | 1 vs 3 | | 0.6189 |
| | | | | | 1 vs 4 | | 0.3455 |
| | | | | | 2 vs 3 | | 0.9927 |
| | | | | | 2 vs 4 | | 0.0486 |
| | | | | | 3 vs 4 | | 0.0327 |
| Fig. 2B | Rotarod | Latency to fall | Two-way RM ANOVA | 12,12,11 | Genotype | F (2, 32) = 56.43 | <0.0001 |
| | 6wks Day 4 | | Tukey's multiple comparisons test | | $Atxn1^{2Q/2Q}$ vs. $Atxn1^{175Q/2Q}$ | | <0.0001 |
| | | | | | $Atxn1^{2Q/2Q}$ vs. $Atxn1^{175QK772T/2Q}$ | | 0.0002 |
| | | | | | $Atxn1^{175Q/2Q}$ vs. $Atxn1^{175QK772T/2Q}$ | | 0.8777 |
| | 12wks Day 4 | | | | $Atxn1^{2Q/2Q}$ vs. $Atxn1^{175Q/2Q}$ | | <0.0001 |
| | | | | | $Atxn1^{2Q/2Q}$ vs. $Atxn1^{175QK772T/2Q}$ | | 0.0007 |
| | | | | | $Atxn1^{175Q/2Q}$ vs. $Atxn1^{175QK772T/2Q}$ | | 0.0327 |
| | 18wks Day 4 | | | | $Atxn1^{2Q/2Q}$ vs. $Atxn1^{175Q/2Q}$ | | <0.0001 |
| | | | | | $Atxn1^{2Q/2Q}$ vs. $Atxn1^{175QK772T/2Q}$ | | 0.0039 |
| | | | | | $Atxn1^{175Q/2Q}$ vs. $Atxn1^{175QK772T/2Q}$ | | <0.0001 |
| | 24wks Day 4 | | | | $Atxn1^{2Q/2Q}$ vs. $Atxn1^{175Q/2Q}$ | | <0.0001 |
| | | | | | $Atxn1^{2Q/2Q}$ vs. $Atxn1^{175QK772T/2Q}$ | | <0.0001 |
| | | | | | $Atxn1^{175Q/2Q}$ vs. $Atxn1^{175QK772T/2Q}$ | | <0.0001 |
| | Rotarod | Latency to fall | Two-way RM ANOVA | 12,12,11 | Age | F (2.518, 80.59) = 14.51 | <0.0001 |
| | $Atxn1^{2Q/2Q}$ | | Tukey's multiple comparisons test | | 6wks vs. 12wks | | 0.7994 |
| | | | | | 6wks vs. 18wks | | 0.6447 |
| | | | | | 6wks vs. 24wks | | 0.4298 |
| | | | | | 12wks vs. 18wks | | 0.9485 |
| | | | | | 12wks vs. 24wks | | 0.7703 |
| | | | | | 18wks vs. 24wks | | 0.9642 |
| | $Atxn1^{175QK772T/2Q}$ | | | | 6wks vs. 12wks | | 0.6173 |
| | | | | | 6wks vs. 18wks | | 0.9980 |
| | | | | | 6wks vs. 24wks | | 0.5243 |
| | | | | | 12wks vs. 18wks | | 0.6031 |
| | | | | | 12wks vs. 24wks | | 0.1924 |
| | | | | | 18wks vs. 24wks | | 0.5516 |
| | $Atxn1^{175Q/2Q}$ | | | | 6wks vs. 12wks | | 0.2584 |
| | | | | | 6wks vs. 18wks | | 0.0167 |
| | | | | | 6wks vs. 24wks | | 0.0002 |
| | | | | | 12wks vs. 18wks | | 0.1130 |
| | | | | | 12wks vs. 24wks | | 0.0001 |
| | | | | | 18wks vs. 24wks | | <0.0001 |
| Fig. 2C | Survival | Lifespan | Log-rank Mantel-Cox test | 22,16 | $Atxn1^{175Q/2Q}$ vs. $Atxn1^{175QK772T/2Q}$ | $X^2$ = 48.12 df = 1 | <0.0001 |
| | | | Gehan-Breslow-Wilcoxon text | | | $X^2$ = 41.11 df = 1 | <0.0001 |
| Fig. 2D | Body Weight | Mass | Two-way RM ANOVA | 12,12,11 | Genotype | F (2, 32) = 23.06 | <0.0001 |
| | 6wks | | Tukey's multiple comparisons test | | $Atxn1^{2Q/2Q}$ vs. $Atxn1^{175Q/2Q}$ | | 0.1020 |
| | | | | | $Atxn1^{2Q/2Q}$ vs. $Atxn1^{175QK772T/2Q}$ | | 0.9423 |
| | | | | | $Atxn1^{175Q/2Q}$ vs. $Atxn1^{175QK772T/2Q}$ | | 0.3057 |
| | 12wks | | | | $Atxn1^{2Q/2Q}$ vs. $Atxn1^{175Q/2Q}$ | | <0.0001 |
| | | | | | $Atxn1^{2Q/2Q}$ vs. $Atxn1^{175QK772T/2Q}$ | | 0.0636 |
| | | | | | $Atxn1^{175Q/2Q}$ vs. $Atxn1^{175QK772T/2Q}$ | | 0.0640 |
| | 18wks | | | | $Atxn1^{2Q/2Q}$ vs. $Atxn1^{175Q/2Q}$ | | <0.0001 |
| | | | | | $Atxn1^{2Q/2Q}$ vs. $Atxn1^{175QK772T/2Q}$ | | 0.0057 |
| | | | | | $Atxn1^{175Q/2Q}$ vs. $Atxn1^{175QK772T/2Q}$ | | 0.0014 |
| | 24wks | | | | $Atxn1^{2Q/2Q}$ vs. $Atxn1^{175Q/2Q}$ | | <0.0001 |
| | | | | | $Atxn1^{2Q/2Q}$ vs. $Atxn1^{175QK772T/2Q}$ | | <0.0001 |
| | | | | | $Atxn1^{175Q/2Q}$ vs. $Atxn1^{175QK772T/2Q}$ | | <0.0001 |
| | Body Weight | Mass | Two-way RM ANOVA | 12,12,11 | Age | F (1.623, 51.94) = 97.10 | <0.0001 |
| | $Atxn1^{2Q/2Q}$ | | Tukey's multiple comparisons test | | 6wks vs. 12wks | | <0.0001 |
| | | | | | 6wks vs. 18wks | | <0.0001 |
| | | | | | 6wks vs. 24wks | | <0.0001 |
| | | | | | 12wks vs. 18wks | | 0.0007 |
| | | | | | 12wks vs. 24wks | | <0.0001 |
| | | | | | 18wks vs. 24wks | | <0.0001 |
| | $Atxn1^{175QK772T/2Q}$ | | | | 6wks vs. 12wks | | <0.0001 |
| | | | | | 6wks vs. 18wks | | <0.0001 |
| | | | | | 6wks vs. 24wks | | <0.0001 |
| | | | | | 12wks vs. 18wks | | 0.1641 |
| | | | | | 12wks vs. 24wks | | 0.7299 |
| | | | | | 18wks vs. 24wks | | 0.5883 |
| | $Atxn1^{175Q/2Q}$ | | | | 6wks vs. 12wks | | 0.0261 |
| | | | | | 6wks vs. 18wks | | 0.6944 |
| | | | | | 6wks vs. 24wks | | 0.3318 |
| | | | | | 12wks vs. 18wks | | 0.1030 |
| | | | | | 12wks vs. 24wks | | 0.0096 |
| | | | | | 18wks vs. 24wks | | 0.0605 |
| Fig. 2E | Brain Weight - 23-26wks | Mass | One-way ANOVA | 4,4,4 | Genotype | F (2, 9) = 31.61 | <0.0001 |
| | | | Tukey's multiple comparisons test | | $Atxn1^{2Q/2Q}$ vs. $Atxn1^{175Q/2Q}$ | | <0.0001 |
| | | | | | $Atxn1^{2Q/2Q}$ vs. $Atxn1^{175QK772T/2Q}$ | | 0.0059 |
| | | | | | $Atxn1^{175Q/2Q}$ vs. $Atxn1^{175QK772T/2Q}$ | | 0.0114 |
| Fig. 3B | Barnes Maze - 7wks | Cognitive score | Two-way RM ANOVA | 10,10,10 | Genotype | F (2, 27) = 4.273 | 0.0244 |
| | Trial 1 | | Tukey's multiple comparisons test | | $Atxn1^{2Q/2Q}$ vs. $Atxn1^{175Q/2Q}$ | | 0.1168 |
| | | | | | $Atxn1^{2Q/2Q}$ vs. $Atxn1^{175QK772T/2Q}$ | | 0.6348 |
| | | | | | $Atxn1^{175Q/2Q}$ vs. $Atxn1^{175QK772T/2Q}$ | | 0.0251 |
| | Trial 2 | | | | $Atxn1^{2Q/2Q}$ vs. $Atxn1^{175Q/2Q}$ | | 0.2403 |
| | | | | | $Atxn1^{2Q/2Q}$ vs. $Atxn1^{175QK772T/2Q}$ | | 0.5889 |
| | | | | | $Atxn1^{175Q/2Q}$ vs. $Atxn1^{175QK772T/2Q}$ | | 0.2976 |
| | Trial 3 | | | | $Atxn1^{2Q/2Q}$ vs. $Atxn1^{175Q/2Q}$ | | 0.6013 |
| | | | | | $Atxn1^{2Q/2Q}$ vs. $Atxn1^{175QK772T/2Q}$ | | 0.9389 |
| | | | | | $Atxn1^{175Q/2Q}$ vs. $Atxn1^{175QK772T/2Q}$ | | 0.7058 |
| | Trial 4 | | | | $Atxn1^{2Q/2Q}$ vs. $Atxn1^{175Q/2Q}$ | | 0.0086 |
| | | | | | $Atxn1^{2Q/2Q}$ vs. $Atxn1^{175QK772T/2Q}$ | | 0.0207 |
| | | | | | $Atxn1^{175Q/2Q}$ vs. $Atxn1^{175QK772T/2Q}$ | | 0.6981 |
| | Barnes Maze - 7wks | Cognitive score | Two-way RM ANOVA | 10,10,10 | Trial Day | F (2.760, 74.51) = 35.81 | <0.0001 |
| | $Atxn1^{2Q/2Q}$ | | Tukey's multiple comparisons test | | 1 vs 2 | | 0.0169 |
| | | | | | 1 vs 3 | | 0.0050 |
| | | | | | 1 vs 4 | | 0.0001 |
| | | | | | 2 vs 3 | | >0.9999 |

| | | | | | | | |
|---|---|---|---|---|---|---|---|
| | | | | | 2 vs 4 | | 0.0228 |
| | | | | | 3 vs 4 | | 0.0228 |
| | $Atxn1^{175QK772T/2Q}$ | | | | 1 vs 2 | | <0.0001 |
| | | | | | 1 vs 3 | | 0.0007 |
| | | | | | 1 vs 4 | | 0.0003 |
| | | | | | 2 vs 3 | | 0.6119 |
| | | | | | 2 vs 4 | | 0.1612 |
| | | | | | 3 vs 4 | | 0.5227 |
| | $Atxn1^{175Q/2Q}$ | | | | 1 vs 2 | | 0.8902 |
| | | | | | 1 vs 3 | | 0.1945 |
| | | | | | 1 vs 4 | | 0.0889 |
| | | | | | 2 vs 3 | | 0.5832 |
| | | | | | 2 vs 4 | | 0.3085 |
| | | | | | 3 vs 4 | | 0.8766 |
| **Fig. 3C** | Barnes Maze - 7wks Trial Day 4 | Cognitive score | One-way ANOVA | 10,10,10 | Genotype | F (2, 27) = 8.730 | 0.0012 |
| | | | Tukey's multiple comparisons test | | $Atxn1^{2Q/2Q}$ vs. $Atxn1^{175Q/2Q}$ | | 0.0016 |
| | | | | | $Atxn1^{2Q/2Q}$ vs. $Atxn1^{175QK772T/2Q}$ | | 0.0084 |
| | | | | | $Atxn1^{175Q/2Q}$ vs. $Atxn1^{175QK772T/2Q}$ | | 0.7940 |
| **Fig. 3D** | Barnes Maze - 17wks | Cognitive score | Two-way RM ANOVA | 8,9,10 | Genotype | F (2, 24) = 41.97 | <0.0001 |
| | Trial 1 | | Tukey's multiple comparisons test | | $Atxn1^{2Q/2Q}$ vs. $Atxn1^{175Q/2Q}$ | | <0.0001 |
| | | | | | $Atxn1^{2Q/2Q}$ vs. $Atxn1^{175QK772T/2Q}$ | | 0.0003 |
| | | | | | $Atxn1^{175Q/2Q}$ vs. $Atxn1^{175QK772T/2Q}$ | | 0.3609 |
| | Trial 2 | | | | $Atxn1^{2Q/2Q}$ vs. $Atxn1^{175Q/2Q}$ | | 0.0073 |
| | | | | | $Atxn1^{2Q/2Q}$ vs. $Atxn1^{175QK772T/2Q}$ | | 0.0578 |
| | | | | | $Atxn1^{175Q/2Q}$ vs. $Atxn1^{175QK772T/2Q}$ | | 0.1536 |
| | Trial 3 | | | | $Atxn1^{2Q/2Q}$ vs. $Atxn1^{175Q/2Q}$ | | 0.0109 |
| | | | | | $Atxn1^{2Q/2Q}$ vs. $Atxn1^{175QK772T/2Q}$ | | 0.1162 |
| | | | | | $Atxn1^{175Q/2Q}$ vs. $Atxn1^{175QK772T/2Q}$ | | 0.0013 |
| | Trial 4 | | | | $Atxn1^{2Q/2Q}$ vs. $Atxn1^{175Q/2Q}$ | | 0.0033 |
| | | | | | $Atxn1^{2Q/2Q}$ vs. $Atxn1^{175QK772T/2Q}$ | | 0.1926 |
| | | | | | $Atxn1^{175Q/2Q}$ vs. $Atxn1^{175QK772T/2Q}$ | | 0.0011 |
| | Barnes Maze - 17wks | Cognitive score | Two-way RM ANOVA | 8,9,10 | Trial Day | F (2.470, 59.29) = 2.950 | 0.0494 |
| | $Atxn1^{2Q/2Q}$ | | Tukey's multiple comparisons test | | 1 vs 2 | | >0.9999 |
| | | | | | 1 vs 3 | | 0.9130 |
| | | | | | 1 vs 4 | | 0.9947 |
| | | | | | 2 vs 3 | | 0.8033 |
| | | | | | 2 vs 4 | | 0.9492 |
| | | | | | 3 vs 4 | | 0.9044 |
| | $Atxn1^{175QK772T/2Q}$ | | | | 1 vs 2 | | 0.1169 |
| | | | | | 1 vs 3 | | 0.0116 |
| | | | | | 1 vs 4 | | 0.0295 |
| | | | | | 2 vs 3 | | 0.3385 |
| | | | | | 2 vs 4 | | 0.0980 |
| | | | | | 3 vs 4 | | >0.9999 |
| | $Atxn1^{175Q/2Q}$ | | | | 1 vs 2 | | 0.9299 |
| | | | | | 1 vs 3 | | 0.4475 |
| | | | | | 1 vs 4 | | 0.9989 |
| | | | | | 2 vs 3 | | 0.8865 |
| | | | | | 2 vs 4 | | 0.9351 |
| | | | | | 3 vs 4 | | 0.0937 |
| **Fig. 3E** | Barnes Maze - 17wks Trial Day 4 | Cognitive score | One-way ANOVA | 8,9,10 | Genotype | F (2, 24) = 16.44 | <0.0001 |
| | | | Tukey's multiple comparisons test | | $Atxn1^{2Q/2Q}$ vs. $Atxn1^{175Q/2Q}$ | | <0.0001 |
| | | | | | $Atxn1^{2Q/2Q}$ vs. $Atxn1^{175QK772T/2Q}$ | | 0.0673 |
| | | | | | $Atxn1^{175Q/2Q}$ vs. $Atxn1^{175QK772T/2Q}$ | | 0.0070 |
| **Fig. 3B-D** | Barnes Maze | $Atxn1^{2Q/2Q}$ cognitive score retention | Paired t test | 8,8 | 7wk Day 4 vs 17wk Day 1 | t=0.4272, df=7 | 0.6821 |
| | | $Atxn1^{175QK772T/2Q}$ cognitive score retention | Paired t test | 9,9 | 7wk Day 4 vs 17wk Day 1 | t=5.330, df=8 | 0.0007 |
| | | $Atxn1^{175Q/2Q}$ cognitive score retention | Paired t test | 10,10 | 7wk Day 4 vs 17wk Day 1 | t=2.744, df=9 | 0.0227 |
| **Fig. 4C** | Subcellular Fractionation - Cerebellum | Expanded ATXN1 nuclear proportion | One-way ANOVA | 4,4,6 | Genotype & age | F (2, 11) = 44.36 | <0.0001 |
| | | | Dunnett's multiple comparisons test | | $Atxn1^{175QK772T/2Q}$ 5wks vs. $Atxn1^{175Q/2Q}$ 5wks | | <0.0001 |
| | | | | | $Atxn1^{175QK772T/2Q}$ 5wks vs. $Atxn1^{175QK772T/2Q}$ 40wks | | 0.0089 |
| **Fig. 4D** | Subcellular Fractionation - Cortex | Expanded ATXN1 nuclear proportion | One-way ANOVA | 4,4,4 | Genotype & age | F (2, 9) = 119.0 | <0.0001 |
| | | | Dunnett's multiple comparisons test | | $Atxn1^{175QK772T/2Q}$ 5wks vs. $Atxn1^{175Q/2Q}$ 5wks | | <0.0001 |
| | | | | | $Atxn1^{175QK772T/2Q}$ 5wks vs. $Atxn1^{175QK772T/2Q}$ 42wks | | 0.0756 |
| **Fig. 4E** | Subcellular Fractionation - Hippocampus | Expanded ATXN1 nuclear proportion | One-way ANOVA | 4,4,4 | Genotype & age | F (2, 9) = 96.27 | <0.0001 |
| | | | Dunnett's multiple comparisons test | | $Atxn1^{175QK772T/2Q}$ 5wks vs. $Atxn1^{175Q/2Q}$ 5wks | | <0.0001 |
| | | | | | $Atxn1^{175QK772T/2Q}$ 5wks vs. $Atxn1^{175QK772T/2Q}$ 42wks | | 0.3001 |
| **Fig. 4F** | Subcellular Fractionation - Medulla | Expanded ATXN1 nuclear proportion | One-way ANOVA | 4,4,6 | Genotype & age | F (2, 11) = 49.01 | <0.0001 |
| | | | Dunnett's multiple comparisons test | | $Atxn1^{175QK772T/2Q}$ 5wks vs. $Atxn1^{175Q/2Q}$ 5wks | | <0.0001 |
| | | | | | $Atxn1^{175QK772T/2Q}$ 5wks vs. $Atxn1^{175QK772T/2Q}$ 40wks | | 0.6982 |
| **Fig. 4G** | Subcellular Fractionation - Striatum | Expanded ATXN1 nuclear proportion | One-way ANOVA | 4,4,4 | Genotype & age | F (2, 9) = 44.70 | <0.0001 |
| | | | Dunnett's multiple comparisons test | | $Atxn1^{175QK772T/2Q}$ 5wks vs. $Atxn1^{175Q/2Q}$ 5wks | | <0.0001 |
| | | | | | $Atxn1^{175QK772T/2Q}$ 5wks vs. $Atxn1^{175QK772T/2Q}$ 42wks | | 0.6593 |
| **Fig. 5A** | Western Blot - Cerebellum | Expanded ATXN1 extractability | Two-way ANOVA | 3,3,3,3 | Genotype | F (1, 8) = 62.49 | <0.0001 |
| | | | | | Age | F (1, 8) = 169.3 | <0.0001 |
| | | | Tukey's multiple comparisons test | | $Atxn1^{175Q/2Q}$ 5wks vs. $Atxn1^{175Q/2Q}$ 26wks | | 0.0023 |
| | | | | | $Atxn1^{175Q/2Q}$ 5wks vs. $Atxn1^{175QK772T/2Q}$ 5wks | | <0.0001 |
| | | | | | $Atxn1^{175Q/2Q}$ 5wks vs. $Atxn1^{175QK772T/2Q}$ 26wks | | 0.0283 |
| | | | | | $Atxn1^{175Q/2Q}$ 26wks vs. $Atxn1^{175QK772T/2Q}$ 5wks | | <0.0001 |
| | | | | | $Atxn1^{175Q/2Q}$ 26wks vs. $Atxn1^{175QK772T/2Q}$ 26wks | | <0.0001 |
| | | | | | $Atxn1^{175QK772T/2Q}$ 5wks vs. $Atxn1^{175QK772T/2Q}$ 26wks | | 0.0023 |
| **Fig. 5D** | Western Blot - Medulla | Expanded ATXN1 extractability | Two-way ANOVA | 3,3,3,3 | Genotype | F (1, 8) = 171.1 | <0.0001 |
| | | | | | Age | F (1, 8) = 24.27 | 0.0012 |
| | | | Tukey's multiple comparisons test | | $Atxn1^{175Q/2Q}$ 5wks vs. $Atxn1^{175Q/2Q}$ 26wks | | 0.1045 |
| | | | | | $Atxn1^{175Q/2Q}$ 5wks vs. $Atxn1^{175QK772T/2Q}$ 5wks | | <0.0001 |
| | | | | | $Atxn1^{175Q/2Q}$ 5wks vs. $Atxn1^{175QK772T/2Q}$ 26wks | | 0.0019 |
| | | | | | $Atxn1^{175Q/2Q}$ 26wks vs. $Atxn1^{175QK772T/2Q}$ 5wks | | <0.0001 |
| | | | | | $Atxn1^{175Q/2Q}$ 26wks vs. $Atxn1^{175QK772T/2Q}$ 26wks | | 0.0001 |
| | | | | | $Atxn1^{175QK772T/2Q}$ 5wks vs. $Atxn1^{175QK772T/2Q}$ 26wks | | 0.0114 |
| **Fig. 5G** | Extractability vs. Subcellular Fractionation Comparison | | Simple linear regression | 4 xy pairs | ATXN1[175QK772T] extractability vs ATXN1[175QK772T] cytoplasmic proportion | $R^2$ = 0.9169 | 0.0425 |
| **Fig. 6G** | Immunofluorescence - Purkinje Cells 21wks | Nuclear Inclusion Frequency | Unpaired t test | 3,3 (98,111) | $Atxn1^{175Q/2Q}$ vs. $Atxn1^{175QK772T/2Q}$ | t=24.99, df=4 | <0.0001 |
| **Fig. 6H** | Immunofluorescence - Medulla 12wks | Nuclear Inclusion Frequency | Unpaired t test | 3,3 (142,137) | $Atxn1^{175Q/2Q}$ vs. $Atxn1^{175QK772T/2Q}$ | t=2.867, df=4 | 0.0456 |
| **Fig. 6I** | Immunofluorescence - Cortex 5wks | Nuclear Inclusion Frequency | Unpaired t test | 3,3 (261,240) | $Atxn1^{175Q/2Q}$ vs. $Atxn1^{175QK772T/2Q}$ | t=4.558, df=4 | 0.0104 |
| **Fig. 6J** | Immunofluorescence - Hippocampus CA1 5wks | Nuclear Inclusion Frequency | Unpaired t test | 3,3 (265,255) | $Atxn1^{175Q/2Q}$ vs. $Atxn1^{175QK772T/2Q}$ | t=34.31, df=4 | <0.0001 |
| **Fig. 6K** | Immunofluorescence - Striatum 5wks | Nuclear Inclusion Frequency | Unpaired t test | 3,3 (281,248) | $Atxn1^{175Q/2Q}$ vs. $Atxn1^{175QK772T/2Q}$ | t=4.128, df=4 | 0.0145 |
| **Fig. 6L** | Immunofluorescence - Cortex 12wks | Nuclear Inclusion Frequency | Unpaired t test | 3,3 (187,313) | $Atxn1^{175Q/2Q}$ vs. $Atxn1^{175QK772T/2Q}$ | t=0.6234, df=4 | 0.5668 |
| **Fig. 6M** | Immunofluorescence - Hippocampus CA1 12wks | Nuclear Inclusion Frequency | Unpaired t test | 3,3 (689,546) | $Atxn1^{175Q/2Q}$ vs. $Atxn1^{175QK772T/2Q}$ | t=0.2868, df=4 | 0.7885 |
| **Fig. 6N** | Immunofluorescence - Striatum 12wks | Nuclear Inclusion Frequency | Unpaired t test | 3,3 (909,528) | $Atxn1^{175Q/2Q}$ vs. $Atxn1^{175QK772T/2Q}$ | t=0.6982, df=4 | 0.5235 |
| **Fig. S1A** | qPCR - Cerebellar Lysates 12wks | Calb1 relative expression | One-way ANOVA | 5,5,3 | Genotype | F (2, 10) = 5.427 | 0.0254 |
| | | | Dunnett's multiple comparisons test | | $Atxn1^{2Q/2Q}$ vs. $Atxn1^{175QK772T/2Q}$ | | 0.0844 |
| | | | | | $Atxn1^{2Q/2Q}$ vs. $Atxn1^{175Q/2Q}$ | | 0.0196 |
| | | Pcp4 relative expression | One-way ANOVA | 5,5,3 | Genotype | F (2, 10) = 24.56 | 0.0001 |
| | | | Dunnett's multiple comparisons test | | $Atxn1^{2Q/2Q}$ vs. $Atxn1^{175QK772T/2Q}$ | | 0.0021 |
| | | | | | $Atxn1^{2Q/2Q}$ vs. $Atxn1^{175Q/2Q}$ | | <0.0001 |
| | | Garnl3 relative expression | One-way ANOVA | 5,5,3 | Genotype | F (2, 10) = 27.73 | <0.0001 |
| | | | Dunnett's multiple comparisons test | | $Atxn1^{2Q/2Q}$ vs. $Atxn1^{175QK772T/2Q}$ | | 0.7550 |
| | | | | | $Atxn1^{2Q/2Q}$ vs. $Atxn1^{175Q/2Q}$ | | <0.0001 |
| | | Rgs8 relative expression | One-way ANOVA | 5,5,3 | Genotype | F (2, 10) = 17.80 | 0.0005 |
| | | | Dunnett's multiple comparisons test | | $Atxn1^{2Q/2Q}$ vs. $Atxn1^{175QK772T/2Q}$ | | 0.4597 |
| | | | | | $Atxn1^{2Q/2Q}$ vs. $Atxn1^{175Q/2Q}$ | | 0.0004 |
| | | Homer3 relative expression | One-way ANOVA | 5,5,3 | Genotype | F (2, 10) = 16.13 | 0.0007 |
| | | | Dunnett's multiple comparisons test | | $Atxn1^{2Q/2Q}$ vs. $Atxn1^{175QK772T/2Q}$ | | 0.0700 |
| | | | | | $Atxn1^{2Q/2Q}$ vs. $Atxn1^{175Q/2Q}$ | | 0.0088 |
| | | Inpp5a relative expression | One-way ANOVA | 5,5,3 | Genotype | F (2, 10) = 7.298 | 0.0111 |
| | | | Dunnett's multiple comparisons test | | $Atxn1^{2Q/2Q}$ vs. $Atxn1^{175QK772T/2Q}$ | | 0.6725 |
| | | | | | $Atxn1^{2Q/2Q}$ vs. $Atxn1^{175Q/2Q}$ | | 0.0076 |
| | | Itpr1 relative expression | One-way ANOVA | 5,5,3 | Genotype | F (2, 10) = 7.430 | 0.0105 |
| | | | Dunnett's multiple comparisons test | | $Atxn1^{2Q/2Q}$ vs. $Atxn1^{175QK772T/2Q}$ | | 0.8499 |
| | | | | | $Atxn1^{2Q/2Q}$ vs. $Atxn1^{175Q/2Q}$ | | 0.0166 |
| **Fig. S1B** | qPCR - Cerebellar Lysates 18wks | Calb1 relative expression | One-way ANOVA | 5,5,5 | Genotype | F (2, 12) = 46.75 | <0.0001 |
| | | | Dunnett's multiple comparisons test | | $Atxn1^{2Q/2Q}$ vs. $Atxn1^{175QK772T/2Q}$ | | <0.0001 |
| | | | | | $Atxn1^{2Q/2Q}$ vs. $Atxn1^{175Q/2Q}$ | | <0.0001 |
| | | Pcp4 relative expression | One-way ANOVA | 5,5,5 | Genotype | F (2, 12) = 20.54 | 0.0001 |
| | | | Dunnett's multiple comparisons test | | $Atxn1^{2Q/2Q}$ vs. $Atxn1^{175QK772T/2Q}$ | | 0.0005 |
| | | | | | $Atxn1^{2Q/2Q}$ vs. $Atxn1^{175Q/2Q}$ | | 0.0001 |
| | | Garnl3 relative expression | One-way ANOVA | 5,5,5 | Genotype | F (2, 12) = 5.259 | 0.0229 |
| | | | Dunnett's multiple comparisons test | | $Atxn1^{2Q/2Q}$ vs. $Atxn1^{175QK772T/2Q}$ | | 0.1531 |
| | | | | | $Atxn1^{2Q/2Q}$ vs. $Atxn1^{175Q/2Q}$ | | 0.0133 |
| | | Rgs8 relative expression | One-way ANOVA | 5,5,5 | Genotype | F (2, 12) = 12.96 | 0.0010 |
| | | | Dunnett's multiple comparisons test | | $Atxn1^{2Q/2Q}$ vs. $Atxn1^{175QK772T/2Q}$ | | 0.0075 |
| | | | | | $Atxn1^{2Q/2Q}$ vs. $Atxn1^{175Q/2Q}$ | | 0.0007 |
| | | Homer3 relative expression | One-way ANOVA | 5,5,5 | Genotype | F (2, 12) = 7.208 | 0.0088 |
| | | | Dunnett's multiple comparisons test | | $Atxn1^{2Q/2Q}$ vs. $Atxn1^{175QK772T/2Q}$ | | 0.0406 |
| | | | | | $Atxn1^{2Q/2Q}$ vs. $Atxn1^{175Q/2Q}$ | | 0.0058 |
| | | Inpp5a relative expression | One-way ANOVA | 5,5,5 | Genotype | F (2, 12) = 10.66 | 0.0022 |
| | | | Dunnett's multiple comparisons test | | $Atxn1^{2Q/2Q}$ vs. $Atxn1^{175QK772T/2Q}$ | | 0.0063 |

| | | | | | | | |
|---|---|---|---|---|---|---|---|
| | | | | | $Atxn1^{2Q/2Q}$ vs. $Atxn1^{175Q/2Q}$ | | 0.0020 |
| | | Itpr1 relative expression | One-way ANOVA | 5,5,5 | Genotype | F (2, 12) = 10.99 | 0.0019 |
| | | | Dunnett's multiple comparisons test | | $Atxn1^{2Q/2Q}$ vs. $Atxn1^{175QK772T/2Q}$ | | 0.0074 |
| | | | | | $Atxn1^{2Q/2Q}$ vs. $Atxn1^{175Q/2Q}$ | | 0.0016 |
| **Fig. S1C** | qPCR - Cerebellar Lysates 26wks | Calb1 relative expression | One-way ANOVA | 4,4,4 | Genotype | F (2, 9) = 43.06 | <0.0001 |
| | | | Dunnett's multiple comparisons test | | $Atxn1^{2Q/2Q}$ vs. $Atxn1^{175QK772T/2Q}$ | | <0.0001 |
| | | | | | $Atxn1^{2Q/2Q}$ vs. $Atxn1^{175Q/2Q}$ | | <0.0001 |
| | | Pcp4 relative expression | One-way ANOVA | 4,4,4 | Genotype | F (2, 9) = 7.471 | 0.0122 |
| | | | Dunnett's multiple comparisons test | | $Atxn1^{2Q/2Q}$ vs. $Atxn1^{175QK772T/2Q}$ | | 0.0280 |
| | | | | | $Atxn1^{2Q/2Q}$ vs. $Atxn1^{175Q/2Q}$ | | 0.0101 |
| | | Garnl3 relative expression | One-way ANOVA | 4,4,4 | Genotype | F (2, 9) = 18.94 | 0.0006 |
| | | | Dunnett's multiple comparisons test | | $Atxn1^{2Q/2Q}$ vs. $Atxn1^{175QK772T/2Q}$ | | 0.0033 |
| | | | | | $Atxn1^{2Q/2Q}$ vs. $Atxn1^{175Q/2Q}$ | | 0.0004 |
| | | Rgs8 relative expression | One-way ANOVA | 4,4,4 | Genotype | F (2, 9) = 47.56 | <0.0001 |
| | | | Dunnett's multiple comparisons test | | $Atxn1^{2Q/2Q}$ vs. $Atxn1^{175QK772T/2Q}$ | | <0.0001 |
| | | | | | $Atxn1^{2Q/2Q}$ vs. $Atxn1^{175Q/2Q}$ | | <0.0001 |
| | | Homer3 relative expression | One-way ANOVA | 4,4,4 | Genotype | F (2, 9) = 43.04 | <0.0001 |
| | | | Dunnett's multiple comparisons test | | $Atxn1^{2Q/2Q}$ vs. $Atxn1^{175QK772T/2Q}$ | | 0.0002 |
| | | | | | $Atxn1^{2Q/2Q}$ vs. $Atxn1^{175Q/2Q}$ | | <0.0001 |
| | | Inpp5a relative expression | One-way ANOVA | 4,4,4 | Genotype | F (2, 9) = 49.60 | <0.0001 |
| | | | Dunnett's multiple comparisons test | | $Atxn1^{2Q/2Q}$ vs. $Atxn1^{175QK772T/2Q}$ | | <0.0001 |
| | | | | | $Atxn1^{2Q/2Q}$ vs. $Atxn1^{175Q/2Q}$ | | <0.0001 |
| | | Itpr1 relative expression | One-way ANOVA | 4,4,4 | Genotype | F (2, 9) = 21.39 | 0.0004 |
| | | | Dunnett's multiple comparisons test | | $Atxn1^{2Q/2Q}$ vs. $Atxn1^{175QK772T/2Q}$ | | 0.0008 |
| | | | | | $Atxn1^{2Q/2Q}$ vs. $Atxn1^{175Q/2Q}$ | | 0.0004 |
| **Fig. S2** | Fear Conditioning - 8wks | % time freezing | One-way ANOVA | 10,10,10 | Genotype | F (2, 27) = 6.597 | 0.0046 |
| | | | Tukey's multiple comparisons test | | $Atxn1^{2Q/2Q}$ vs. $Atxn1^{175Q/2Q}$ | | 0.0100 |
| | | | | | $Atxn1^{2Q/2Q}$ vs. $Atxn1^{175QK772T/2Q}$ | | 0.9975 |
| | | | | | $Atxn1^{175Q/2Q}$ vs. $Atxn1^{175QK772T/2Q}$ | | 0.0118 |
| **Fig. S4A** | Western Blot - Cortex | Expanded ATXN1 extractability | Two-way ANOVA | 3,3,3,3 | Genotype | F (1, 8) = 894.4 | <0.0001 |
| | | | | | Age | F (1, 8) = 24.51 | 0.0011 |
| | | | Tukey's multiple comparisons test | | $Atxn1^{175Q/2Q}$ 5wks vs. $Atxn1^{175Q/2Q}$ 26wks | | 0.5016 |
| | | | | | $Atxn1^{175Q/2Q}$ 5wks vs. $Atxn1^{175QK772T/2Q}$ 5wks | | <0.0001 |
| | | | | | $Atxn1^{175Q/2Q}$ 5wks vs. $Atxn1^{175QK772T/2Q}$ 26wks | | <0.0001 |
| | | | | | $Atxn1^{175Q/2Q}$ 26wks vs. $Atxn1^{175QK772T/2Q}$ 5wks | | <0.0001 |
| | | | | | $Atxn1^{175Q/2Q}$ 26wks vs. $Atxn1^{175QK772T/2Q}$ 26wks | | <0.0001 |
| | | | | | $Atxn1^{175QK772T/2Q}$ 5wks vs. $Atxn1^{175QK772T/2Q}$ 26wks | | 0.0024 |
| **Fig. S4D** | Western Blot - Hippocampus | Expanded ATXN1 extractability | Two-way ANOVA | 3,3,3,3 | Genotype | F (1, 8) = 1391 | <0.0001 |
| | | | | | Age | F (1, 8) = 212.9 | <0.0001 |
| | | | Tukey's multiple comparisons test | | $Atxn1^{175Q/2Q}$ 5wks vs. $Atxn1^{175Q/2Q}$ 26wks | | 0.0861 |
| | | | | | $Atxn1^{175Q/2Q}$ 5wks vs. $Atxn1^{175QK772T/2Q}$ 5wks | | <0.0001 |
| | | | | | $Atxn1^{175Q/2Q}$ 5wks vs. $Atxn1^{175QK772T/2Q}$ 26wks | | <0.0001 |
| | | | | | $Atxn1^{175Q/2Q}$ 26wks vs. $Atxn1^{175QK772T/2Q}$ 5wks | | <0.0001 |
| | | | | | $Atxn1^{175Q/2Q}$ 26wks vs. $Atxn1^{175QK772T/2Q}$ 26wks | | <0.0001 |
| | | | | | $Atxn1^{175QK772T/2Q}$ 5wks vs. $Atxn1^{175QK772T/2Q}$ 26wks | | <0.0001 |
| **Fig. S4G** | Western Blot - Striatum | Expanded ATXN1 extractability | Two-way ANOVA | 3,3,3,3 | Genotype | F (1, 8) = 153.6 | <0.0001 |
| | | | | | Age | F (1, 8) = 15.99 | 0.0040 |
| | | | Tukey's multiple comparisons test | | $Atxn1^{175Q/2Q}$ 5wks vs. $Atxn1^{175Q/2Q}$ 26wks | | 0.7918 |
| | | | | | $Atxn1^{175Q/2Q}$ 5wks vs. $Atxn1^{175QK772T/2Q}$ 5wks | | <0.0001 |
| | | | | | $Atxn1^{175Q/2Q}$ 5wks vs. $Atxn1^{175QK772T/2Q}$ 26wks | | 0.0016 |
| | | | | | $Atxn1^{175Q/2Q}$ 26wks vs. $Atxn1^{175QK772T/2Q}$ 5wks | | <0.0001 |
| | | | | | $Atxn1^{175Q/2Q}$ 26wks vs. $Atxn1^{175QK772T/2Q}$ 26wks | | 0.0006 |
| | | | | | $Atxn1^{175QK772T/2Q}$ 5wks vs. $Atxn1^{175QK772T/2Q}$ 26wks | | 0.0065 |
| **Fig. S5A** | Extractability - Cerebellum 5wks | Total expanded ATXN1 | Unpaired t test | 3,3 | $Atxn1^{175Q/2Q}$ vs. $Atxn1^{175QK772T/2Q}$ | t=2.631, df=4 | 0.0581 |
| **Fig. S5B** | Subcellular Fractionation - Cerebellum 5wks | Total expanded ATXN1 | Unpaired t test | 4,4 | $Atxn1^{175Q/2Q}$ vs. $Atxn1^{175QK772T/2Q}$ | t=2.151, df=6 | 0.0750 |
| **Fig. S5C** | Extractability - Cortex 5wks | Total expanded ATXN1 | Unpaired t test | 3,3 | $Atxn1^{175Q/2Q}$ vs. $Atxn1^{175QK772T/2Q}$ | t=15.08, df=4 | 0.0001 |
| **Fig. S5D** | Subcellular Fractionation - Cortex 5wks | Total expanded ATXN1 | Unpaired t test | 4,4 | $Atxn1^{175Q/2Q}$ vs. $Atxn1^{175QK772T/2Q}$ | t=16.91, df=6 | <0.0001 |
| **Fig. S5E** | Extractability - Hippocampus 5wks | Total expanded ATXN1 | Unpaired t test | 3,3 | $Atxn1^{175Q/2Q}$ vs. $Atxn1^{175QK772T/2Q}$ | t=23.26, df=4 | <0.0001 |
| **Fig. S5F** | Subcellular Fractionation - Hippocampus 5wks | Total expanded ATXN1 | Unpaired t test | 4,4 | $Atxn1^{175Q/2Q}$ vs. $Atxn1^{175QK772T/2Q}$ | t=6.768, df=6 | 0.0005 |
| **Fig. S5G** | Extractability - Medulla 5wks | Total expanded ATXN1 | Unpaired t test | 3,3 | $Atxn1^{175Q/2Q}$ vs. $Atxn1^{175QK772T/2Q}$ | t=2.966, df=4 | 0.0413 |
| **Fig. S5H** | Subcellular Fractionation - Medulla 5wks | Total expanded ATXN1 | Unpaired t test | 4,4 | $Atxn1^{175Q/2Q}$ vs. $Atxn1^{175QK772T/2Q}$ | t=6.007, df=6 | 0.0010 |
| **Fig. S5I** | Extractability - Striatum 5wks | Total expanded ATXN1 | Unpaired t test | 3,3 | $Atxn1^{175Q/2Q}$ vs. $Atxn1^{175QK772T/2Q}$ | t=8.783, df=4 | 0.0009 |
| **Fig. S5J** | Subcellular Fractionation - Striatum 5wks | Total expanded ATXN1 | Unpaired t test | 4,4 | $Atxn1^{175Q/2Q}$ vs. $Atxn1^{175QK772T/2Q}$ | t=21.22, df=6 | <0.0001 |

**Table 1. Detailed statistical summary of results**

| Primer | Sequence 5'->3' | Probe Source | Probe Name | Probe Catalogue # |
|---|---|---|---|---|
| mCalb1 712F | AAGGCTTTTGAGTTATATGATCAGG | Roche | 42 | UPL 04688015001 |
| mCalb1 800R | TTCTTCTCACACAGATCTTTCAGC | Roche | | |
| mPcp4 205F | CCAACGGAAAAGACAAGACG | Roche | 29 | UPL 04687612001 |
| mPcp4 276R | TGTCGATATCAAATTCTTCTTGGA | Roche | | |
| mGarnl3 1516F | TCATGAAGCCGTGTGTGC | Roche | 89 | UPL 04689143001 |
| mGarnl3 1609R | CACGGATGGGAGGTCATC | Roche | | |
| mRgs8 561F | CTGTCACACAAATCAGACTCCTG | Roche | 88 | UPL 04689135001 |
| mRgs8 653R | TGCTTCTTCCGTGGAGAGTC | Roche | | |
| mHomer3 682F | TGAAGAAGATGCTGTCAGAAGG | Roche | 13 | UPL 04685121001 |
| mHomer3 754R | CTGTCCTGAAGCGCGAAG | Roche | | |
| mInpp5a 688F | ATTCGGACACTTTGGAGAGC | Roche | 67 | UPL 04688660001 |
| mInpp5a 775R | CCTTTTCTTGACCATTTGCAC | Roche | | |
| mItpr1 13462F | GAAGGCATCTTTGGAGGAAGT | Roche | 85 | UPL 04689097001 |
| mItpr1 13538R | ACCCTGAGGAAGGTTCTGC | Roche | | |
| mGapdh | Roche Proprietery Information | Roche | - | 5046211001 |
| mAtxn1 F | CACGGTCATTCAGACCACACA | IDT | Custom Probe | FAM-CAGCCACGG/ZEN/CCTTCTACGCTGG/3IABKFQ/ |
| mAtxn1 R | GGTAGCCGATGACAGGAGGTT | IDT | | |

**Table S2. List of oligonucleotides and probes used for RT-qPCR in this study**
